# Glia detect and mount a protective response to loss of dendrite substructure integrity in *C. elegans*

**DOI:** 10.1101/2023.11.16.567404

**Authors:** Katherine C. Varandas, Brianna M. Hodges, Lauren Lubeck, Amelia Farinas, Yupu Liang, Yun Lu, Shai Shaham

## Abstract

Neurons have elaborate structures that determine their connectivity and functions. Changes in neuronal structure accompany learning and memory formation and are hallmarks of neurological disease. Here we show that glia monitor dendrite structure and respond to dendrite perturbation. In *C. elegans* mutants with defective sensory-organ dendrite cilia, adjacent glia accumulate extracellular matrix-laden vesicles, secrete excess matrix around cilia, alter gene expression, and change their secreted protein repertoire. Inducible cilia disruption reveals that this response is acute. DGS-1, a 7-transmembrane domain neuronal protein, and FIG-1, a multifunctional thrombospondin-domain glial protein, are required for glial detection of cilia integrity, and exhibit mutually-dependent localization to and around cilia, respectively. While inhibiting glial secretion disrupts dendritic cilia properties, hyperactivating the glial response protects against dendrite damage. Our studies uncover a homeostatic protective dendrite-glia interaction and suggest that similar signaling occurs at other sensory structures and at synapses, which resemble sensory organs in architecture and molecules.

## INTRODUCTION

The elaborate structures of neurons determine connectivity and functional output in the nervous system. Neurons typically extend axon and dendrite polarized projections that mediate signal transmission and reception, respectively.^1^ Many axons are ensheathed by glia, some of which can form densely layered myelinating sheets.^2^ Likewise, dendritic spines, which mediate reception of presynaptically-released neurotransmitters,^3^ are often surrounded by astrocyte processes,^4^ and microtubule-rich cilia at dendritic endings of sensory neurons are also enveloped by adjacent glia.^5^

While glia surrounding axons are known to detect axon diameter for proportional myelinatation^6^ and respond to axon damage to promote or block nerve regeneration and repair,^7^ glial responses to dendrite structural perturbations are far less understood. Yet, changes in dendritic substructures are commonplace during learning and memory formation, and are a hallmark of many nervous system diseases.^8^ For example, ciliopathies, genetic disorders leading to abnormal cilia structure, often cause profound sensory deficits.^9^ We set out, therefore, to determine whether glia can monitor and respond to changes in dendrite structure.

The sensory organs of *C. elegans* are an excellent setting to study dendrite-glia communication. The majority of *C. elegans* glia physically associate with dendritic endings at sensory organs^10^ and these interactions are invariant to a resolution of tens of nanometers.^11–13^ Cilia are only found at sensory dendrite endings in *C. elegans*,^14^ allowing us to model loss of dendrite substructure using cilia mutants. The amphids are a pair of bilaterally-symmetric head sensory organs, each containing twelve sensory neurons, eight of which project axons into the brain-like nerve ring and extend anterior dendrites ending in simple cilia housing sensory receptors and signal transduction machinery.^11^ These cilia are required for sensory neuron function,^15^ and protrude through a channel formed by two glial cells: the amphid sheath (AMsh) glial cell, which secretes an extracellular matrix around the cilia, and the amphid socket (AMso) glial cell, which forms a pore through which cilia are exposed to the environment.^11,16^ Four other amphid neurons have elaborately-structured dendritic endings that are ensheathed only by AMsh glia: three with elaborately-structured wing cilia dendrite endings, and another with actin-rich microvilli dendrite ending (**Figure 1A**).^17,18^

**Figure 1.**
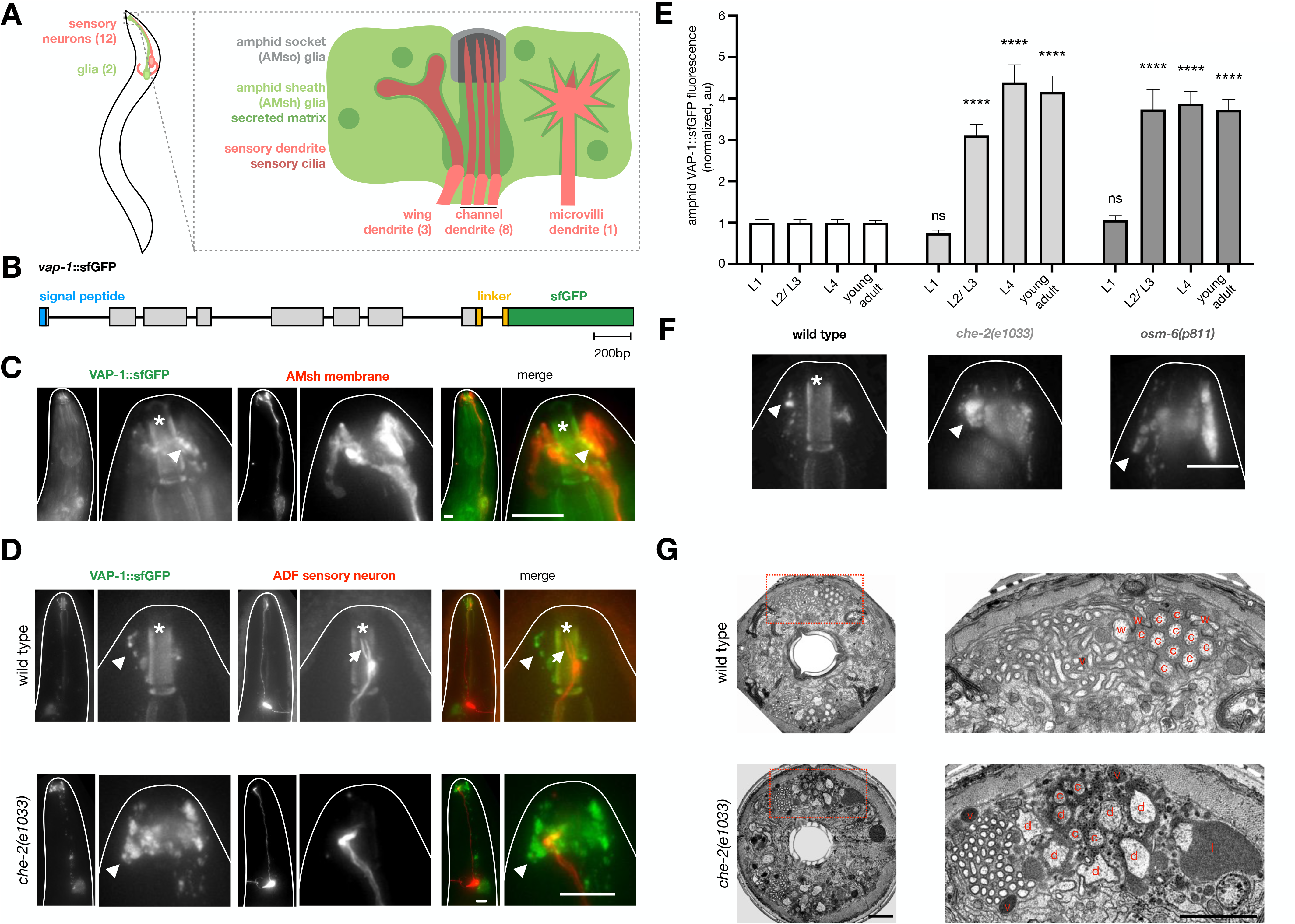
AMsh glia modify their secretory activity in dendrite cilia mutants. (**A**) Schematic of an amphid sensory organ. (**B**) CRISPR edited *vap-1*::*sfGFP* locus. (**C**) *vap-1::sfGFP* animal expressing AMsh glia membrane reporter (*F16F9.3pro::myr-mKate2*). GFP channel is background subtracted. Arrowheads, VAP-1::sfGFP puncta. Asterisk, anterior buccal cavity auto-fluorescence. Scale bar, 10μm. (**D**) Representative brightness and contrast-matched images of wild type and *che-2(e1033)* animals co-expressing *vap-1::sfGFP* and the ADF neuron reporter (*srh-142pro::dsRed*). Arrows, dendrite cilia. Arrowheads, VAP-1::sfGFP puncta. Asterisk, anterior buccal cavity auto-fluorescence. Scale bar=10μm. (**E**) Quantification of VAP-1::sfGFP fluorescence in wild-type, *che-2(e1033)*, and *osm-6(p811)* amphids over larval development. Statistical significance with respect to wild type animals at the same timepoint, calculated using Dunnett’s multiple comparison test following 2-way ANOVA. *n*=29-33 animals/condition from 3 independent experiments. Error bars, SEM. ns, p>0.05. ****, p<0.0001. (**F**) Representative brightness and contrast matched images of VAP-1::sfGFP in young adult animals of indicated genotypes. Arrowheads, asterisk, as in (C). Scale bar=10μm. (**G**) Full TEM cross sections (left) and insets (right) of wild type and *che-2(e1033)* mutant animals. c, channel cilia. w, wing cilia. d, dendrite. L, amphid channel lobe. v, matrix-filled AMsh vesicle. Scale bars=2μm. All fluorescence images are maximum intensity projections of widefield z-stacks.

Here we report that AMsh glia actively monitor the structural integrity of sensory dendrite cilia and respond protectively when ciliary defects are detected. In mutants with defective cilia, AMsh glia secrete excess matrix around dendrite endings, and concomitantly accumulate matrix-laden secretory vesicles. These changes in secretion dynamics are accompanied by transcriptional changes in the AMsh glial cell that alter the secretory repertoire of the cell and correlate with increased activity of the conserved glial transcription factor PROS-1. By inducibly disrupting cilia, we show that the glial secretory response is acute. We identify two genes, *dgs-1* and *fig-1*, that mediate glial detection of and responses to cilia changes. *dgs-1* encodes a previously uncharacterized 7-transmembrane domain protein expressed in a subset of amphid sensory neurons and localized to neuronal cilia. *fig-1* encodes a transmembrane thrombospondin-domain protein expressed in AMsh glia and localized around amphid channel cilia. DGS-1 and FIG-1 protein localization is mutually dependent. In either *dgs-1* or *fig-1* mutants, cilia are intact, yet AMsh glia exhibit secretory changes similar to those seen in mutants with abnormal cilia structure. Mutation of *dgs-1* delays the onset of cilia dysfunction following inducible cilia disruption, revealing that the glial responses are protective. Our studies uncover and highlight the importance of glia in monitoring and maintaining homeostasis of signaling compartments in the nervous system.

## RESULTS

### AMsh glia modify their secretory activity in dendrite cilia mutants

To determine whether glia monitor and respond to loss of dendrite substructure integrity, we examined whether disruption of *C. elegans* sensory dendrite cilia affects surrounding AMsh glia. AMsh glia express many secreted proteins,^19^ raising the possibility that cilia disruption could affect glial secretion. To test this idea, we first generated a reporter to monitor AMsh glia secretory activity. We used CRISPR/Cas9 to generate animals in which the endogenous *vap-1* gene, encoding an AMsh glia-enriched secreted protein,^19^ is fused in frame to coding sequences for superfolder GFP (sfGFP), optimized for fluorescence in the extracellular milieu (**Figure 1B**).^20^ In these animals, VAP-1::sfGFP localizes to discreet puncta surrounding sensory neuron cilia at the anterior tip of AMsh glia, as expected for a protein packaged into secretory vesicles and then released around sensory cilia (**Figures 1C and 1D**, top panel).

We next examined VAP-1::sfGFP expression *che-2(e1033)* mutant animals, which have truncated cilia.^18,21^ *che-2* encodes a homolog of the IFT80 subunit of the ciliary intraflagellar transport B (IFT-B) complex and is expressed exclusively in ciliated sensory neurons in *C. elegans*.^14,21^ We observed increased VAP-1::sfGFP accumulation around amphid sensory cilia in these mutants (**Figure 1D**, compare top and bottom panels). Similar accumulation was found in loss-of-function mutants of the ciliary rootlet gene *che-10*/rootletin, the IFT-A complex gene *che-11*/IFT140, and the IFT-B complex genes *osm-6*/IFT52, *osm-5*/IFT88 and *dyf-11*/IFT54 (**Figure S1A**), all of which exhibit truncated cilia.^21–23^ To determine when VAP-1::sfGFP accumulation begins, we imaged in developmentally-synchronized *che-2(e1033)* and *osm-6(p811)* mutants at 24-hour intervals over larval development. In both, excess VAP-1::sfGFP accumulation is first evident in L2/L3 larvae and persists through adulthood (**Figures 1E and 1F**).

To confirm these results, we examined the amphid ultrastructure of wild type and *che-2(e1033)* mutant animals using serial-section transmission electron microscopy (TEM; 2 amphids observed for each strain). In *che-2(e1033)* mutants, we observed excess extracellular matrix in lobe-like structures protruding from the AMsh glia channel membrane surrounding dendrite endings, as well as excess vesicles near the glial membrane (**Figure 1G**, compare top and bottom panels). These changes are consistent with our VAP-1::sfGFP findings and previous TEM studies suggesting that AMsh glia accumulate vesicles containing extracellular matrix in cilia mutants.^18,24^

Taken together, our data demonstrate that AMsh glia detect the loss of dendrite cilia integrity and respond by increasing the number of extracellular matrix-laden vesicles around the glial channel housing these cilia, and by secreting excess extracellular matrix into this channel.

### AMsh glia transcription is altered in neuronal cilia mutants and correlates with increased PROS-1/Prox1 activity

To further explore how loss of cilia integrity affects AMsh glia, we performed RNA sequencing (RNA-seq) on cells isolated from either wild type or *che-2(e1033)* mutant late-stage larvae expressing an AMsh glia-specific reporter (*F16F9.3*pro::dsRed). AMsh glia (dsRed+) from each strain were sorted from all other cells (dsRed-) using fluorescence activated cell sorting (FACS) and processed as previously described.^19^ For both wild type animals and *che-2* mutants, comparison of dsRed+ AMsh glia transcripts versus those in dsRed-cells shows enrichment of mRNAs previously described as AMsh glia-enriched, validating our cell isolation and RNA-seq methodology (**Figures S2A and S2B**).^19,25^ This finding also demonstrates that AMsh glia cell fate is not entirely altered in cilia mutants, as at least some AMsh glia genes are enriched to the same degree in wild-type and *che-2(e1033)* animals. Nonetheless, we found several hundred genes whose expression was either increased (673 genes; FC ≥ 2, padj ≤ 0.1) or decreased (485 genes; FC ≤ 0.5, padj ≤ 0.1) in AMsh glia of *che-2* cilia mutants (**Figure S2C**). Strikingly, of the 673 genes with increased expression in *che-2* mutants, 185 (27.59%) are predicted to encode membrane proteins and 24 (3.57%) are predicted to encode secreted proteins. Of the 485 downregulated genes, 159 (32.78%) are predicted to encode membrane proteins and 30 (6.19%) are predicted to encode secreted proteins. Thus, in addition to accumulating secreted matrix, glia respond to defects in dendrite cilia by altering the expression of over 1,000 genes, many of which encode proteins predicted to enter the secretory pathway.

PROS-1, a major transcriptional regulator of the AMsh glia secretome,^19,26^ is homologous to *Drosophila* Prospero and to mammalian Prox1, both of which are also expressed in sensory organ glia.^19,27–29^ Consistent with the idea that the secretory pathway in AMsh glia is altered upon cilia disruption, AMsh glia genes whose expression is induced or repressed by PROS-1 cluster with genes up-regulated or down-regulated in AMsh glia of *che-2(e1033)* mutants, respectively, more than expected statistically (**Figures S2C and S2D**). PROS-1-induced genes are, on average, more enriched in AMsh glia of *che-2(e1033)* mutants than PROS-1-repressed genes (**Figure S2E**). These data suggest that the transcriptional response of AMsh glia to cilia defects may, at least in part, result from increased activation of PROS-1, which controls the secretory repertoire of AMsh glia.

Expression of a PROS-1::GFP reporter is not increased in AMsh glia of cilia mutants and *pros-1* mRNA levels are similarly unaffected (**Figures S2F and S2G**). Furthermore, although VAP-1::sfGFP accumulates in the glial tips of cilia mutants, we do not detect an increase in *vap-1* mRNA in these mutants (**Figure S2H**). Therefore, in addition to transcriptional changes, post-transcriptional changes affecting the secretory pathway must also take place in AMsh glia in response to cilia disruption.

### The AMsh glia response to cilia disruption is acute

Animals carrying mutations in *che-2*, *osm-6*, or other cilia genes have chronic cilia defects, which can be monitored by assaying the ability of a subset of ciliated neurons to take up the small-molecule dyes DiI or DiO following incubation of animals with these fluorophores.^15^ To determine whether the glial response to cilia defects can also follow acute cilia perturbation, we designed an inducible cilia degradation system. Briefly, sequences encoding AID, an <u>a</u>uxin-inducible <u>d</u>egron,^30,31^ were introduced into the genomic *osm-6* locus of animals also ubiquitously expressing the AID-targeting scaffold protein TIR1 (*eft-3pro::TIR1-mRuby*; **Figure 2A**; a related method is described in ^32^). While control animals carrying either *osm-6::AID* or *eft-3pro::TIR1-mRuby*, have no dye-filling defects, we found that in the majority of animals carrying both constructs, sensory neurons become dye-filling defective within 1.5 hours of auxin exposure (**Figure 2B**). TEM micrographs of a dye-filling defective animal reveal loss of cilia integrity (**Figure S3**; 2 amphids observed). Furthermore, *osm-6::GFP-AID* animals, in which we can follow OSM-6 protein, show loss of GFP fluorescence following auxin exposure (**Figure 2C**), confirming OSM-6 protein degradation.

**Figure 2.**
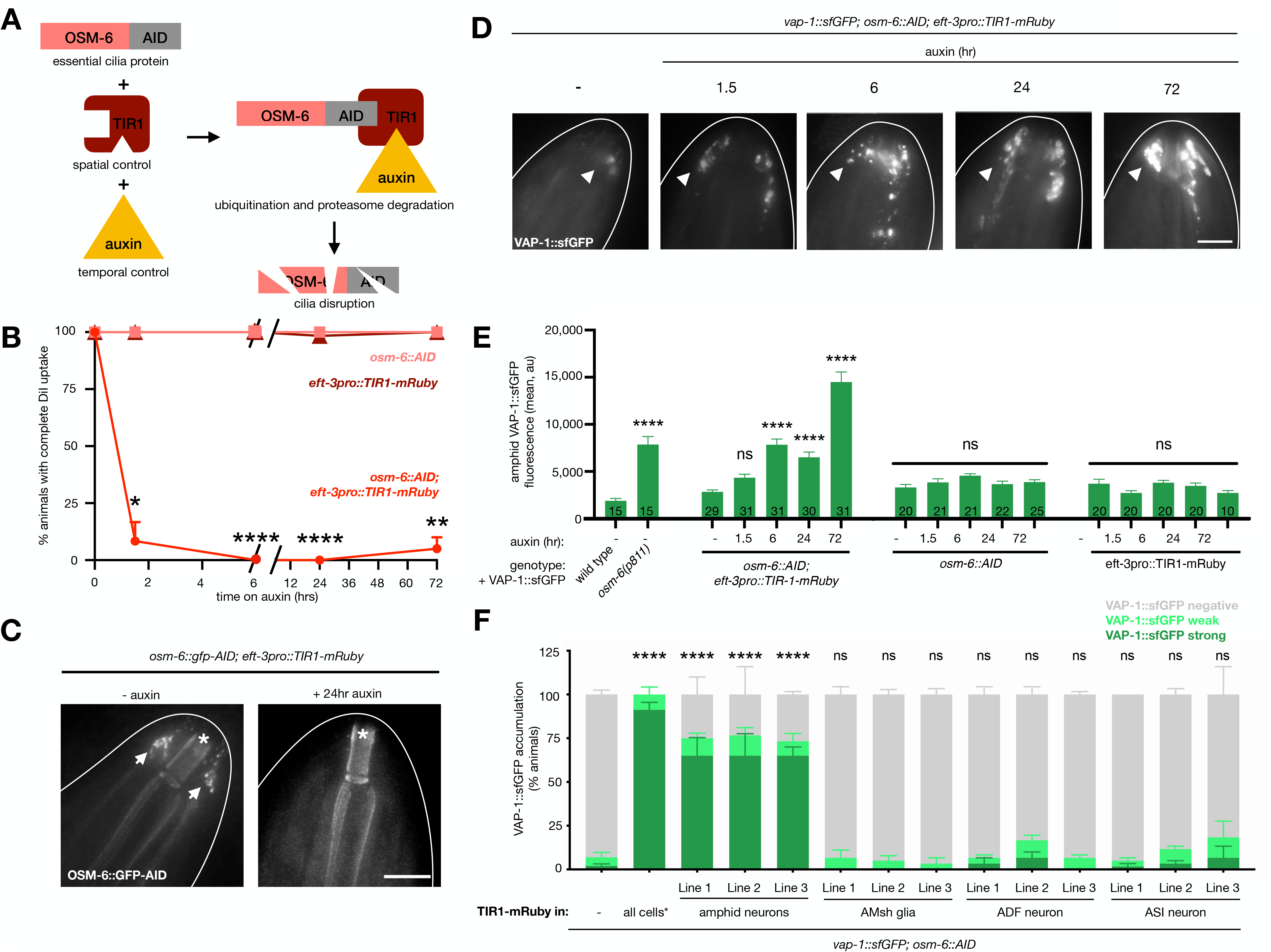
The AMsh glia response to cilia disruption is acute. (**A**) Strategy for inducible cilia disruption. (**B**) Amphid dye-filling time course of animals with indicated inducible cilia disruption components. *n*=3 trials/condition, 20 animals/ trial. Statistical significance shown in reference to same genotype without auxin treatment condition, calculated using Dunnett’s multiple comparison test following 2-way ANOVA. (**C**) Representative brightness and contrast-matched images of *osm-6::gfp-AID* in animals also ubiquitously expressing TIR1-mRuby expression (*eft-3pro::TIR1-mRuby*) without auxin or following 24 hours of auxin treatment. Arrows, OSM-6::GFP-AID in amphid. Asterisks, anterior buccal cavity auto-fluorescence. (**D**) Representative brightness and contrast-matched images of VAP-1::sfGFP in *osm-6::AID* animals also ubiquitously expressing TIR1-mRuby expression (*eft-3pro::TIR1-mRuby*) without auxin or treated with auxin for indicated time. Arrowheads, VAP-1::sfGFP puncta. (**E**) Quantification of VAP-1::sfGFP accumulation in the amphid of animals with indicated genotypes and auxin treatment time. *n* (total animals quantified) indicated inside bars. Statistical significance shown in relation to condition without auxin of the same genotype, or wild type condition for *osm-6(p811)*, calculated using Tukey’s multiple comparison test following one-way ANOVA. ns, p>0.05. ****, p<0.0001. (**F**) VAP-1::sfGFP accumulation in *osm-6::AID* animals expressing TIR1-mRuby in indicated cell types. *n*≥3 trials/ condition, 20 animals scored/trial. Statistical significance shown in relation to animals with no TIR1-mRuby calculated using % animals with strong VAP-1::sfGFP accumulation with Dunnett’s multiple comparison test following one-way ANOVA. *, TIR1-mRuby is expressed in all cells (*eft-3pro::TIR1-mRuby*) from a genetically integrated transgene, while cell-specific TIR1-mRuby is expressed from extrachromosomal arrays. ns, p>0.05. ****, p<0.0001. All images are maximum intensity projections of widefield z-stacks. Scale bars= 10μm. All data are represented as mean+SEM.

We used our inducible cilia degradation system to assess AMsh glia response dynamics following acute cilia disruption. We exposed developmentally-synchronized *vap-1::sfGFP; osm-6::AID; eft-3pro::TIR1-mRuby* animals to auxin at different times prior to imaging as young adults and found that VAP-1::sfGFP progressively accumulates at the AMsh glia tip (**Figures 2D and 2E**). Control animals carrying only *osm-6::AID* or *eft-3pro::TIR1-mRuby*, show no VAP-1::sfGFP accumulation (**Figure 2E**). We conclude that AMsh glia respond within hours to disruption of dendrite cilia, and, therefore, that this response must be mediated by a neuronal signal with similar or faster action.

To determine whether cilia disruption of multiple channel neurons is required to see a glial response, we constructed *vap-1::sfGFP; osm-6::AID* animals that express TIR1-mRuby only in specific cells or cell combinations. When TIR1-mRuby is expressed in all amphid neurons, VAP-1::sfGFP accumulates in most animals exposed to auxin for 72 hours. Animals in which TIR1-mRuby is expressed in AMsh glia show no accumulation (**Figure 2F**), consistent with the notion that OSM-6 acts only in neuronal cilia and indicating that glia are detecting a signal from associated neurons. Importantly, disruption of either ADF neuron or ASI neuron cilia alone results in VAP-1::sfGFP accumulation in only a few animals (**Figure 2F**). Thus, AMsh glia require disruption of multiple dendrite cilia to elicit the full matrix accumulation response.

### *dgs-1* mutants have intact cilia, yet aberrantly accumulate AMsh glia-secreted matrix

To identify genes involved in the AMsh glia response to dendrite cilia loss, we mutagenized *vap-1::sfGFP* animals also expressing an ADF neuron reporter, and screened F2 progeny for mutants that accumulate VAP-1::sfGFP at AMsh glia tips. To exclude mutants with cilia structural defects, only candidates with normal ADF cilia morphology and whose amphid neurons take up DiI were further studied. From this screen, we isolated a homozygous mutant, *ns942*, in which VAP-1::sfGFP accumulation is pronounced despite normal cilia structure (**Figure 3A**).

**Figure 3.**
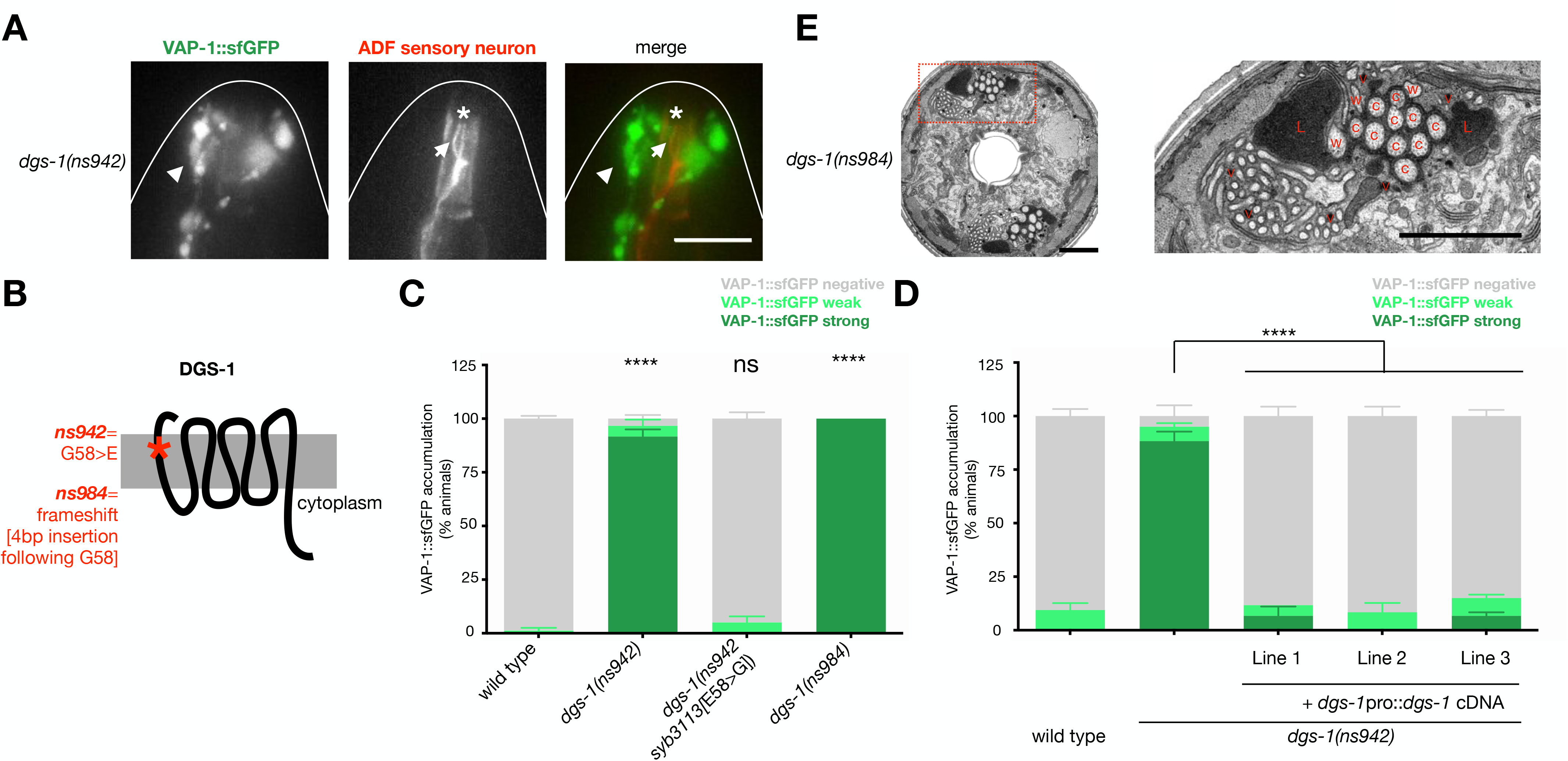
*dgs-1* mutants have intact cilia, yet aberrantly accumulate AMsh glia-secreted matrix. (**A**) Representative widefield images of a *dgs-1(ns942)*; *vap-1::sfGFP* animal expressing an ADF neuron reporter (*srh-142*pro::dsRed). VAP-1::sfGFP images are brightness and contrast-matched to Figure 1D for comparison. Arrows, dendrite cilia. Arrowheads, VAP-1 puncta. Asterisk, anterior buccal cavity auto-fluorescence. Scale bar= 10μm. (**B**) DGS-1 predicted topology and mutations analyzed. (**C**) VAP-1::sfGFP accumulation in animals of indicated genotype. *n*≥3 trials, 20 animals/trial. ns, p>0.05. ****, p<0.0001. (**D**) VAP-1::sfGFP accumulation in animals of indicated genotype. *n*≥3 trials, 20 animals/trial. ns, p>0.05. ****, p<0.0001. (**E**) Full TEM section through a *dgs-1(ns942)*; *vap-1::sfGFP* animal expressing an ADF neuron reporter (*srh-142*pro::dsRed; left). Inset on right. c, channel cilia. w, wing cilia. L, amphid channel lobe. v, matrix-filled AMsh vesicle. Scale bars=2μm. Fluorescence images are maximum intensity projections of widefield z-stacks. Data are represented as mean+SEM. Statistical significance calculated using Tukey’s multiple comparison following one-way ANOVA of % animals with strong VAP-1::sfGFP accumulation and shown in relation to the wild type unless otherwise indicated.

Using whole genome sequencing, single-nucleotide polymorphism mapping, and transformation rescue, we demonstrated that *ns942* is a recessive allele with a causal lesion in a previously uncharacterized gene, which we renamed <u>d</u>endrite-<u>g</u>lia-<u>s</u>ignaling-1 (*dgs-1*). *dgs-1* encodes a predicted 7-transmembrane domain protein, and *ns942* mutants have a predicted glycine-to-glutamic acid substitution in the first transmembrane domain (**Figure 3B**). To confirm that *dgs-1* is the relevant gene, we showed that animals in which the *ns942* lesion is repaired (*dgs-1(ns942 syb3113[E58>G])*) exhibit no VAP-1::sfGFP accumulation (**Figure 3C**). Furthermore, animals homozygous for a loss-of-function frameshift allele we generated (*ns984*), in which four base pairs are inserted following the G58 coding sequence, also exhibit aberrant VAP-1::sfGFP accumulation (**Figure 3C**). Finally, *ns942* mutants expressing a *dgs-1* cDNA using *dgs-1* regulatory sequences (*dgs-1*pro::*dgs-1* cDNA), show near complete rescue of the VAP-1::sfGFP accumulation defect (**Figure 3D**).

To determine whether *dgs-1* mutants have ultrastructural changes similar to those of cilia mutants, we examined *dgs-1(ns984)* animals by TEM. Indeed, while amphid cilia of *dgs-1* mutants appear grossly intact, vesicles filled with electron-dense matrix accumulate in *dgs-1* mutants and the amphid channel membrane has large lobe-like extensions containing electron-dense secreted matrix material (**Figure 3E**; 6 amphids observed). Thus, despite having intact cilia, *dgs-1* mutants exhibit AMsh glia secretion defects resembling those of *che-2* and other cilia mutants.

### DGS-1 functions in sensory neurons to signal the presence of intact dendrite cilia to surrounding glia

To determine where DGS-1 functions, we first examined expression of a *dgs-1pro::GFP* reporter transgene. We observed GFP fluorescence in six of eight amphid channel sensory neurons and in neurons of the tail phasmid sensory organ (**Figures 4A and S4A**), which are similar in structure and gene expression to the amphid.^15^ We identified four of the neurons that express *dgs-1* as ADL, ASK, ASH, and ASJ by co-labeling with DiI (**Figure 4A**). The remaining two neurons, ADF and ASG, were identified by co-labeling with neuron-specific reporters (**Figures S4B** and **S4C**). Supporting these cell assignments, expression data from the CeNGEN database^33^ as well as from our AMsh glia RNA-seq experiments shows that in the amphid, *dgs-1* transcripts are enriched only in the six neurons we identified and not in other neurons or glia (**Figure 4B**). To confirm that DGS-1 functions in sensory neurons, we expressed *dgs-1* cDNA using an amphid sensory neuron-specific promoter, *dyf-11*pro, in *dgs-1(ns942)* mutants, and saw strong rescue of VAP-1::sfGFP accumulation (**Figure 4C**).

**Figure 4.**
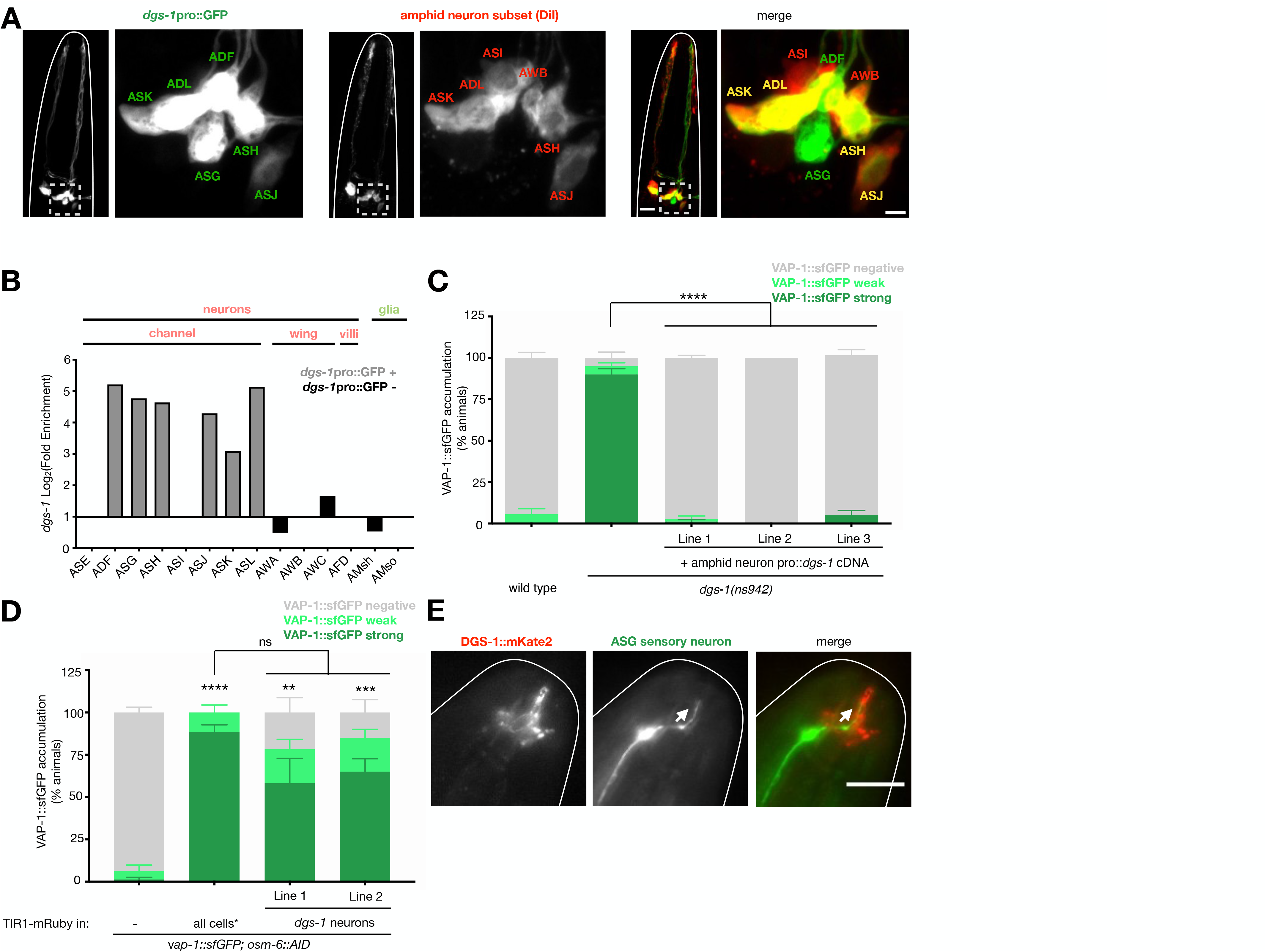
DGS-1 functions in sensory neurons to signal the presence of intact dendrite cilia to surrounding glia. (**A**) Representative confocal images of a *dgs-*1pro::GFP animal labeled with DiI. Left panels, head region. Right panels, inset showing neuron cell bodies of one amphid. Scale bars, head region=10um, inset= 2μm. (**B**) RNA-seq expression of *dgs-1* in the amphid. Cells with no RNA-seq reads have no bars. All data is from the CeNGEN project^33^ except AMsh glia, which is from our RNA-seq experiments (presented in Figure S2). (**C**) VAP-1::sfGFP accumulation in wild type and *dgs-1(ns942)* animals, and in 3 independent lines expressing *dgs-1* cDNA using an amphid neuron promoter (*dyf-11)* in *dgs-1(ns942)*. ns, P>0.05. ****, P≤0.0001. (**D**) VAP-1::sfGFP scoring in *osm-6::AID* animals expressing TIR1-mRuby in all cells or only from the *dgs-1* promoter. *,TIR1-mRuby expressed in all cells (*eft-3pro::TIR1-mRuby*) is expressed from a genetically integrated transgene, while *dgs-1* neuron-specific TIR1-mRuby is expressed from an extrachromosomal array. ns, P>0.05. **, P≤0.01. ***, P≤0.001. ****, P≤0.0001. (**E**) Representative wide-field image of an animal expressing DGS-1::mKate2 and ASG neuron reporter (*ops-1*pro::GFP). Scalebar=10μm. Arrow, dendrite cilia. Images are maximum intensity projections of z-stacks. For VAP-1::sfGFP scoring experiments, *n*≥3 trials/ condition, 20 animals scored/trial. Data are represented as mean+SEM. Statistics calculated using Tukey’s multiple comparison following one-way ANOVA from % animals with strong VAP-1::sfGFP accumulation.

To further confirm the DGS-1 site of action, we expressed TIR1-mRuby using the *dgs-1* promoter in *vap-1::sfGFP; osm-6::AID* animals. Transgenic animals exposed to auxin for 72 hours displayed pronounced VAP-1::sfGFP accumulation, comparable to that seen in animals in which cilia of all amphid neurons are disrupted (**Figure 4D**, compare to Figure 2F). Thus, perturbing only the neurons in which DGS-1 is expressed is sufficient to elicit the full glial response. DGS-1 must, therefore, be a component of the main, or only, neuron-glia signaling interaction reporting on dendrite cilia integrity.

To determine where DGS-1 localizes within sensory neurons, we inserted mKate2 fluorescent protein coding sequences into the predicted third intracellular loop of *dgs-1* (DGS-1::mKate2, **Figure S4D**). This fusion protein is fully functional, as animals expressing the tagged protein in *dgs-1(ns942)* animals have no AMsh glia secretory changes (**Figure S4E**). We find that DGS-1::mKate2 localizes to cilia at the anterior tips of amphid sensory dendrites (**Figure 4E**). Together, our data demonstrate that DGS-1 functions in the cilia of six amphid sensory neurons to signal the presence of intact dendrite cilia to the surrounding AMsh glial cell.

### FIG-1, an AMsh glia-expressed protein, is required for monitoring cilia integrity

While DGS-1 functions in neurons to report on dendrite cilia integrity, glial proteins must exist that receive this neuronal information. We reasoned that such proteins are likely to be transmembrane proteins whose inactivation promotes the glial response. Indeed, we found that deleting the gene *fig-1*, encoding a glia-expressed transmembrane protein with a large extracellular domain,^25^ results in strong VAP-1::sfGFP accumulation (*fig-1*(Δcoding sequence), **Figures 5A and 5B**). Like *dgs-1* and *che-2* mutants, serial-section

**Figure 5.**
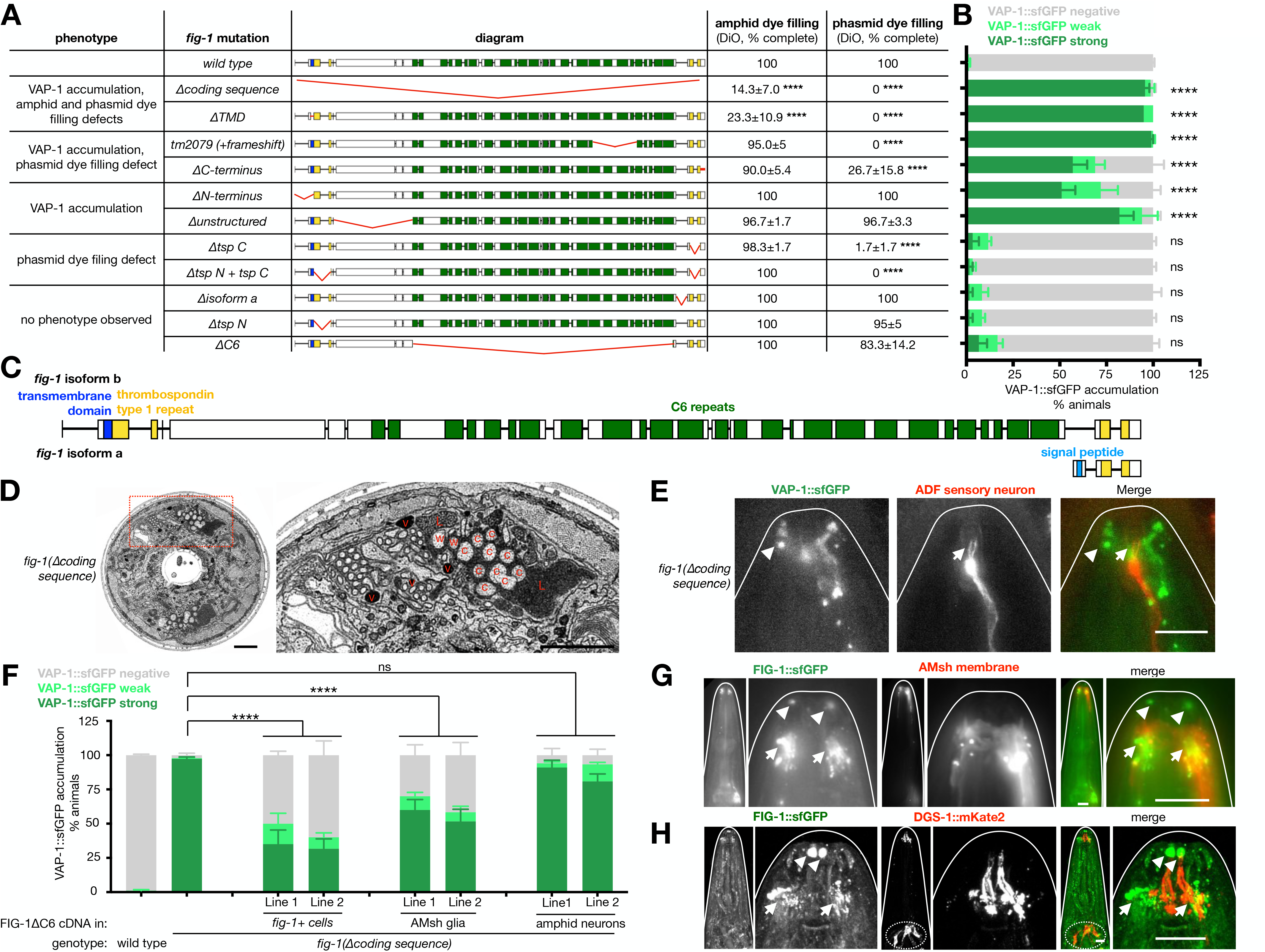
FIG-1, an AMsh glia-expressed protein, is required for monitoring cilia integrity. (**A**) Schematic of *fig-1* deletion mutants and DiO dye filling, grouped by phenotype. Dye filling is indicated as mean±SEM of *n*≥3 trials, 20 animals/per trial. Statistical significance shown relative to wild type, calculated using Dunnett’s multiple comparison following one-way ANOVA, only annotated if significant. ****, P≤0.0001. (**B**) VAP-1::sfGFP accumulation in wild type animals and *fig-1* mutants corresponding to (A). Statistical significance shown in relation to wild type animals, calculated using Dunnett’s multiple comparison following one-way ANOVA from % animals with strong VAP-1::sfGFP accumulation. ns, P>0.05. ****, P≤0.0001. (**C**) *fig-1* predicted gene structure and protein domains. (**D**) Full TEM cross section of *fig-1(Δcoding sequence)* mutant (left), with enlarged inset (right). c, channel cilia. w, wing cilia. L, amphid channel lobes. V, matrix-filled AMsh vesicle. Scale bars=2μm. (**E**) Representative images of *vap-1::sfGFP* animal expressing an ADF neuron reporter (*srh-142pro::dsRed*) in *fig-1(Δcoding sequence)* mutant. Arrows, cilia. Arrowheads, VAP-1::sfGFP puncta. Scale bar= 10μm. (**F**) VAP-1::sfGFP accumulation in wild type, *fig-1(Δcoding sequence)*, and 2 independent lines expressing *fig-1Δ*C6 cDNA from the *fig-1* promoter, an AMsh glia-specific promoter (*F16F9.3*), or an amphid neuron specific promoter (*dyf-11*). Statistical significance calculated using Tukey’s multiple comparison test following one-way ANOVA. ns, P>0.05. ****, P≤0.0001. (**G**) Representative widefield images of a *fig-1::sfGFP* animal expressing AMsh membrane reporter *F16F9.3*pro::myr-mKate2. Arrowheads, anterior FIG-1::sfGFP puncta. Arrows, FIG-1::sfGFP within AMsh glia. Scale bar=10μm. (**H**) Representative confocal images of *fig-1*::sfGFP animal also expressing DGS-1::mKate2. Arrowheads, anterior FIG-1::sfGFP puncta. Arrows, FIG-1::sfGFP around cilia base. Scale bar= 10μm. Fluorescence images are maximum intensity projections z-stacks. Data are represented as mean+SEM.

TEM reveals that this mutant has large amphid channel lobes containing matrix material and also accumulates secretary vesicles carrying matrix near the channel membrane (**Figure 5D**; 2 amphids observed). Similar defects are observed in TEM micrographs of another *fig-1* mutant, *fig-1(tm2079)* (**Figure S5A**; 4 amphids observed, see below). Importantly, sensory neuron cilia appear normal in ultrastructure and by fluorescent imaging in *fig-1*(Δcoding sequence) and *fig-1(tm2079)* mutants (**Figures 5D, 5E, and 6C**), consistent with our previous work showing that these neurons are also largely functional.^25^

**Figure 6.**
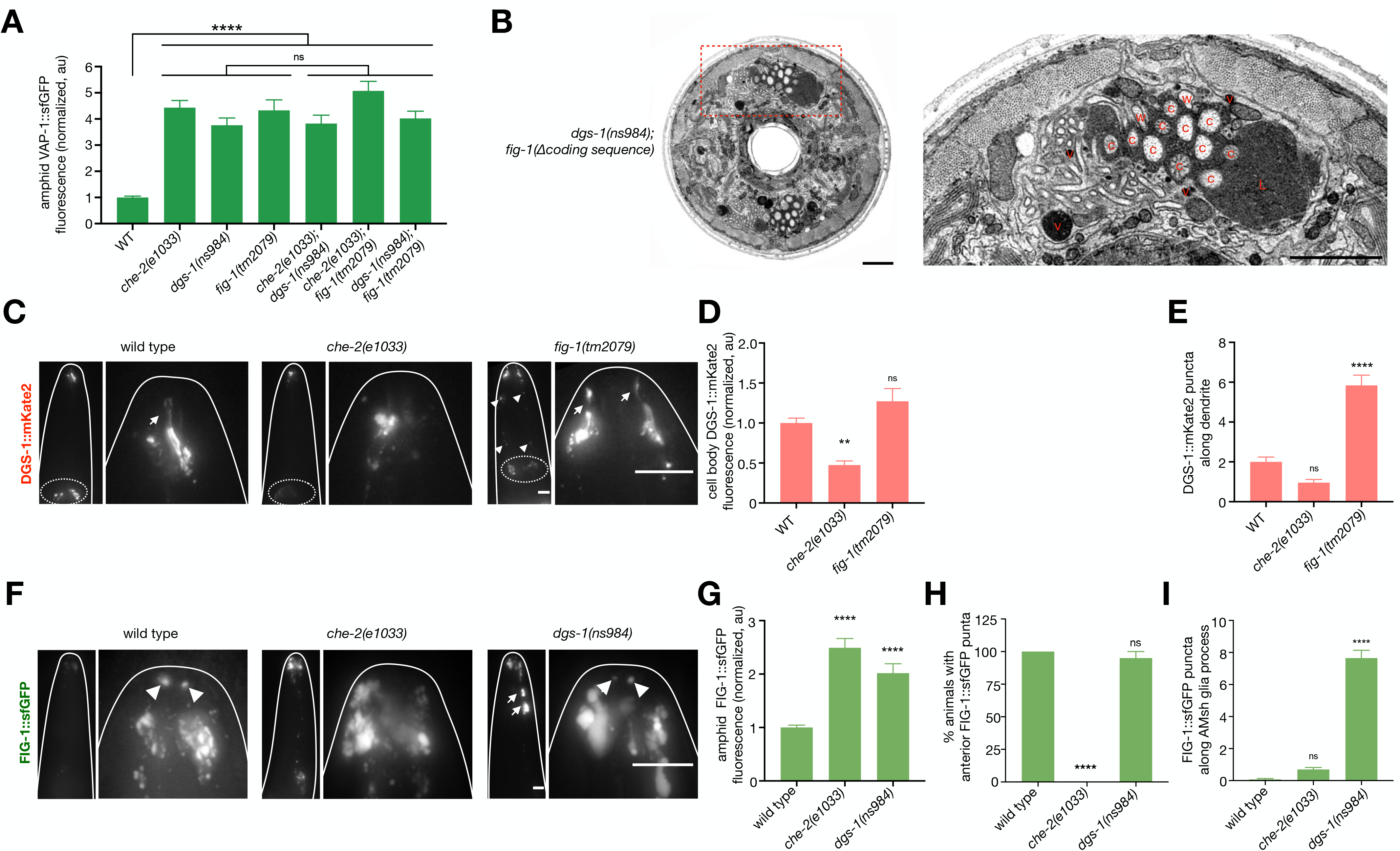
DGS-1 and FIG-1 function in the same pathway. (**A**) VAP-1::sfGFP quantification in the amphids of animals with indicated genotypes. *n*=30 animals from 3 independent experiments. ns, P>0.05. ****, P≤0.0001. (**B**) Full TEM cross section through *dgs-1(ns984)*; *fig-1(Δcoding sequence)* mutant (left), and inset (right). c, channel cilia. w, wing cilia. L, amphid channel lobes. v, matrix-filled AMsh vesicle. Scale bars=2μm. (**C**) Representative brightness and contrast-matched images of animals expressing DGS-1::mKate2 in a wild type animal, and *che-2(e1033)* and *fig-1(tm2079)* mutants. Left, animal head. Right, inset of nose tip. Arrows, dendrite cilia. Arrowheads, DGS-1::mKate2 accumulation along dendrites. Dotted ellipses, neuron cell bodies. Scale bar=10μm. (**D**) Quantification of DGS-1::mKate2 fluorescence in neuron cell bodies. *n*=40 animals from 4 independent experiments. ns, P>0.05. **, P≤0.01. (**E**) Quantification of DGS-1::mKate2 puncta along the dendrites (between buccal cavity and cell bodies) per animal. *n*=40 animals from 4 independent experiments. ns, P>0.05. ****, P≤0.0001. (**F**) Representative images of FIG-1::sfGFP a wild type animal, and *che-2(e1033)* and *dgs-1(ns984)* mutants. Left, brightness and contrast-matched animal head images. Right, inset of nose tip scaled to show FIG-1::sfGFP localization. Arrowheads, anterior FIG-1::sfGFP puncta. Arrows, FIG-1::sfGFP in ≥2um diameter puncta along the AMsh glial processes. Scale bar=10μm. (**G**) Quantification of FIG-1::sfGFP fluorescence in the amphid. *n*=40 animals from 4 independent experiments. ****, P≤0.0001. (**H**) Quantification of percent of animals with at least one anterior FIG-1::sfGFP puncta visible/ image stack. *n*=40 animals from 4 independent experiments. ns, P>0.05. ****, P≤0.0001. (**I**) Quantification of FIG-1::sfGFP puncta ≥2um diameter along the glial processes per animal, defined as between buccal cavity and cell body. *n*=40 animals from 4 independent experiments ns, P>0.05. ****, P≤0.0001. All fluorescence images are maximum intensity projections of widefield z-stacks. Data are represented as mean+SEM. Statistical significance calculated using Tukey’s multiple comparison test following one-way ANOVA and shown in relation to wild type animals unless otherwise indicated.

Unlike *dgs-1* mutants, and despite having structurally normal cilia, *fig-1(Δ coding sequence)* mutants exhibit defects in dye uptake by amphid and phasmid neurons (**Figure 5A**).^25^ These findings suggest that FIG-1 has multiple functions, one monitoring cilia integrity and another facilitating dye uptake. To test this idea, and to identify FIG-1 domains required for these functions, we performed *in vivo* structure-function analysis of the gene. The *fig-1* locus generates two transcripts, predicted to encode proteins of 138 and 2898 amino acids (FIG-1a and FIG-1b, respectively; **Figure 5C**). We found that deletion of the *fig-1* intron containing the first exon of FIG-1a, *fig-1(Δ isoform a)*, has no effect on either VAP-1::sfGFP accumulation or dye uptake (**Figures 5A and 5B**). Thus, FIG-1a appears to have at most a minor role, and we therefore focused our attention on FIG-1b.

FIG-1b protein contains a short N-terminal intracellular region, a transmembrane domain, and two thrombospondin type I (TSP) domains separated by 17 C6 repeats that constitute the bulk of the protein (**Figure 5C**). We found that like the *Δ coding sequence* mutant, in-frame *in vivo* deletion of the transmembrane domain (*fig-1ΔTMD*, amino acids 23-51), perturbs dye filling and also results in VAP-1::sfGFP accumulation (**Figures 5A and 5B**), indicating that this deletion destroys protein function. Deletion of the C-terminal 24 amino acids of FIG-1b (*fig-1ΔC-terminus*, amino acids 2,874-2,898), or deletion of several C6 repeats, followed by a frame-shift (allele *tm2079*, amino acids 2250-2593 and early stop), perturbs dye filling in the phasmid sensory organ and also results in VAP-1::sfGFP accumulation (**Figures 5A and 5B**). We conclude that this 24 amino acid sequence at the C-terminus of the protein, is critical for both functions.

Thrombospondin domain-containing proteins are secreted from mammalian glia and function in synaptogenesis.^34^ We wondered, therefore, what roles the two TSP1 domains of *fig-1* may play. We found that an in-frame deletion of the C-terminal thrombospondin type 1 repeat (*fig-1ΔtspC*, amino acids 2,823-2,873) causes dye-filling defects in the phasmid and has no effect on VAP-1::sfGFP accumulation (**Figure 5A and 5B**). Deletion in-frame of the N-terminal TSP1 domain has little effect on dye filling or VAP-1::sfGFP, and deleting both TSP1 domains does not enhance the defects over what is observed for deletion of the C-terminal TSP1 domain alone (**Figures 5A and 5B**). Thus, the C-terminal TSP1 domain is required for phasmid dye uptake but not for VAP-1::sfGFP accumulation in AMsh glia, suggesting these two functions are independent.

Supporting this idea, deleting in frame the cytoplasmic N-terminal domain (*fig-1ΔN-terminus*, amino acids 2-22) or the unstructured extracellular domain preceding the C6 repeats (*fig-1Δunstructured*, amino acids 115-754; **Figures 5A and 5B**), promotes aberrant VAP-1::sfGFP accumulation, but has no effect on dye uptake in either the amphid or phasmid. Taken together, these findings reveal that FIG-1b is a multifunctional glial protein with separable functions, with some domains required only for VAP-1::sfGFP accumulation, and others only for dye uptake.

Surprisingly, an in-frame deletion spanning all 17 C6 repeats (*fig-1ΔC6*, amino acids 755-2788) results in little detectable VAP-1 accumulation or dye-filling defects (**Figures 5A and 5B**). Additionally, expression of a *fig-1* cDNA lacking the coding region for the C6-repeats under endogenous *fig-1* gene regulatory sequences (*fig-1pro::fig-1*Δ*C6* cDNA) rescues the VAP-1::sfGFP accumulation defect of *fig-1*(Δ coding sequence) mutants (**Figure 5F**), demonstrating that the C6 repeats are indeed largely dispensable for FIG-1b function.

### FIG-1 functions from AMsh glia and localizes near sensory neuron cilia

We previously demonstrated that *fig-1* is expressed only in AMsh and phasmid sheath glia.^25^ To determine in which cell(s) FIG-1 functions to facilitate normal AMsh matrix secretion, we expressed FIG-1b lacking the dispensable C6 repeat region in amphid neurons and AMsh glia of *fig-1(Δcoding sequence)* mutants. We were unable to amplify a full-length *fig-1b* cDNA for this study, as it is large and contains a repeated structure. While little rescue is observed when *fig-1*Δ*C6* is expressed from an amphid neuron-specific promoter, rescue is observed when an AMsh glia promoter is used (**Figure 5F**). Thus, FIG-1 functions in AMsh glia to control AMsh glia matrix secretion.

To determine where in AMsh glia FIG-1 is localized, we generated animals in which sfGFP coding sequences are inserted into the endogenous *fig-1* locus just before the stop codon (FIG-1::sfGFP). Since these animals do not exhibit dye-filling defects (**Figure S5B**), we believe that the FIG-1::sfGFP fusion protein generated is functional. We observed FIG-1::sfGFP expression in AMsh glia, as confirmed by co-expression of the AMsh glia reporter *F16F9.3pro::myr-mKate2* (**Figure 5G**), localized to the anterior part of AMsh glia surrounding the ciliary base (**Figure 5H**). This distribution is consistent with a FIG-1b being a transmembrane protein localized to the amphid channel, and mirrors the localization of VAP-1. We also consistently observed two discreet FIG-1::sfGFP puncta outside of the AMsh glia membrane (**Figure 5G and 5H**, arrowheads), which are located at the tips of amphid channel dendrite cilia marked by DGS-1::mKate2 (**Figure 5H**). These two distinct localization sites may reflect the two different functions of FIG-1. Although FIG-1::sfGFP tags both FIG-1a and FIG-1b isoforms, anterior puncta are also seen when FIG-1 is tagged internally (**Figure S5C**), indicating that these puncta house FIG-1b. These observations, therefore, suggest that FIG-1b protein is targeted to anterior puncta is released from the AMsh glia membrane. Together, our data indicate that FIG-1 is carried in AMsh glia by vesicles resembling those transporting VAP-1, and functions from these glia in reception of a cilia integrity signal.

### DGS-1 and FIG-1 function in the same pathway

To test if *dgs-1*, *fig-1*, and cilia genes such as *che-2*, function in the same pathway, we generated animals carrying pair-wise combinations of mutations in these genes and assayed activation of the glial response. We found that double-mutant animals accumulate VAP-1::sfGFP in the amphid to the same extent as each single mutant (**Figure 6A**). Supporting this conclusion, serial-section TEM of *dgs-1(ns984)*; *fig-1(Δcoding sequence)* double mutants does not reveal additional channel or vesicle-accumulation defects compared to either single mutant (**Figure 6B**, 2 amphids observed; compare to **Figures 3E** and **5D**). One interpretation of these findings is that mutations in *che-2*, *dgs-1* or *fig-1* each activate the glial response to its maximal extent, precluding an enhancement of the response. However, because mutations in the gene *daf-6*, a regulator of amphid channel size, result in matrix accumulation well beyond what we see in these mutants,^20^ we believe that such a ceiling effect is unlikely to explain our double mutant findings. We therefore favor the conclusion that double mutants fail to give an enhanced glial response because all three genes function in the same pathway.

To further probe the relationship between *dgs-1, fig-1,* and dendrite cilia mutants, we examined the DGS-1::mKate2 reporter in cilia and *fig-1* mutants. Although DGS-1::mKate2 is still transported to cilia-disrupted dendrite tips in *che-2(e1033)* mutants (**Figure 6C**, arrows), overall expression of this reporter is greatly reduced (note neuronal cell bodies in **Figures 6C**, dotted circles, **and 6D**). This result is consistent with CHE-2 and DGS-1 functioning in the same pathway, and suggests that activation of the glial response in *che-2(e1033)* mutants may be due to a combination of reduced DGS-1 expression and inappropriate DGS-1 localization at the ciliary base due to cilia disruption. In *fig-1(tm2079)* mutants, DGS-1::mKate2 levels are not significantly changed and the protein localizes to dendrite cilia (**Figures 6C and 6D**). However, the reporter accumulates abnormally in puncta along the dendrites (**Figures 6C**, arrowheads, **and 6E**), indicating that FIG-1 guides or stabilizes DGS-1 localization at the dendrite tip.

We also examined FIG-1::sfGFP localization and expression in *che-2* and *dgs-1* mutants. We observed a marked increase in FIG-1::sfGFP accumulation at the glial tip in both mutants (**Figures 6F**, compare brightness and contrast matched whole head images, **and 6G**). These findings mirror changes we observe in VAP-1 in these mutants (**Figures 1D-1F**, **3A, and 3C**), supporting the notion that FIG-1 travels through the same secretory route as VAP-1. We did not observe an increase in *fig-1* mRNA levels in *che-2(e1033)* mutants, however (**Figure S6**), indicating that changes in FIG-1 abundance must occur post-transcriptionally. In *che-2*, but not *dgs-1*, mutants, we also observed loss of the two anterior FIG-1::sfGFP puncta near amphid dendrite cilia tips (**Figures 6F**, arrowheads, and **Figure 6H**). Importantly, just as DGS-1 aberrantly accumulates in puncta along sensory neuron dendrites in *fig-1* mutants, FIG-1 accumulates in puncta along the AMsh glial process of *dgs-1* mutants (**Figures 6F**, arrowheads, **and 6I**). Thus, DGS-1 reciprocally guides or stabilizes FIG-1 localization.

In summary, our genetic and expression data support the conclusion that cilia genes, *dgs-1*, and *fig-1* function in the same pathway to induce AMsh glia responses to ensheathed dendrite cilia damage. Our data also suggest that *fig-1* functions downstream of cilia and *dgs-1* mutations, as mutations in *fig-1* do not alter the levels of DGS-1 as *che-2* mutations do. Conversely, both *che-2* and *dgs-1* mutations increase accumulation of FIG-1 in the amphid, indicating that they function upstream of FIG-1.

### The AMsh glial response protects neuronal cilia

Previous studies from our lab suggest that secretion from AMsh glia may be important for sensory neuron function. For example, knockdown of *pros-1* results in loss of expression of many secreted and transmembrane proteins, blocks dye uptake by sensory neurons, and perturbs behavior in response to sensory stimuli.^19^ To test this idea directly, we sought to block secretion from AMsh glia by expressing a dominant-negative RAB-1 protein, RAB-1(S25N), in these cells using the AMsh-specific promoter from the *ver-1* gene, which is active at 25C but not 15C.^35^ RAB-1 is a key regulator of endoplasmic-reticulum-to-Golgi transport in a wide range of organisms, and its inactivation blocks cellular secretion.^36^ We previously showed that RAB-1(S25N) expression in AMsh glia disrupts the microvillar structure of AFD neuron dendritic endings.^35^ Maintenance of these endings requires glial membrane accumulation of the KCC-3 transporter, consistent with the idea that RAB-1(S25N) can perturb secretion in AMsh glia.^35^ As shown in **Figure 7A**, while dye uptake is not perturbed in amphid sensory neurons of transgenic *ver-1* promoter::RAB-1(S25N) animals or their non-transgenic siblings raised at 15C, ~15% of *ver-1* promoter::RAB-1(S25N) animals raised at 25C, but not their non-transgenic siblings, fail to uptake DiO, and a similar proportion of animals exhibit weaker dye filling than wild-type animal. This result shows that AMsh glia secretion is required for dye uptake by amphid sensory neurons, and raises the possibility that increasing secretion from AMsh glia could protect damaged cilia.

**Figure 7.**
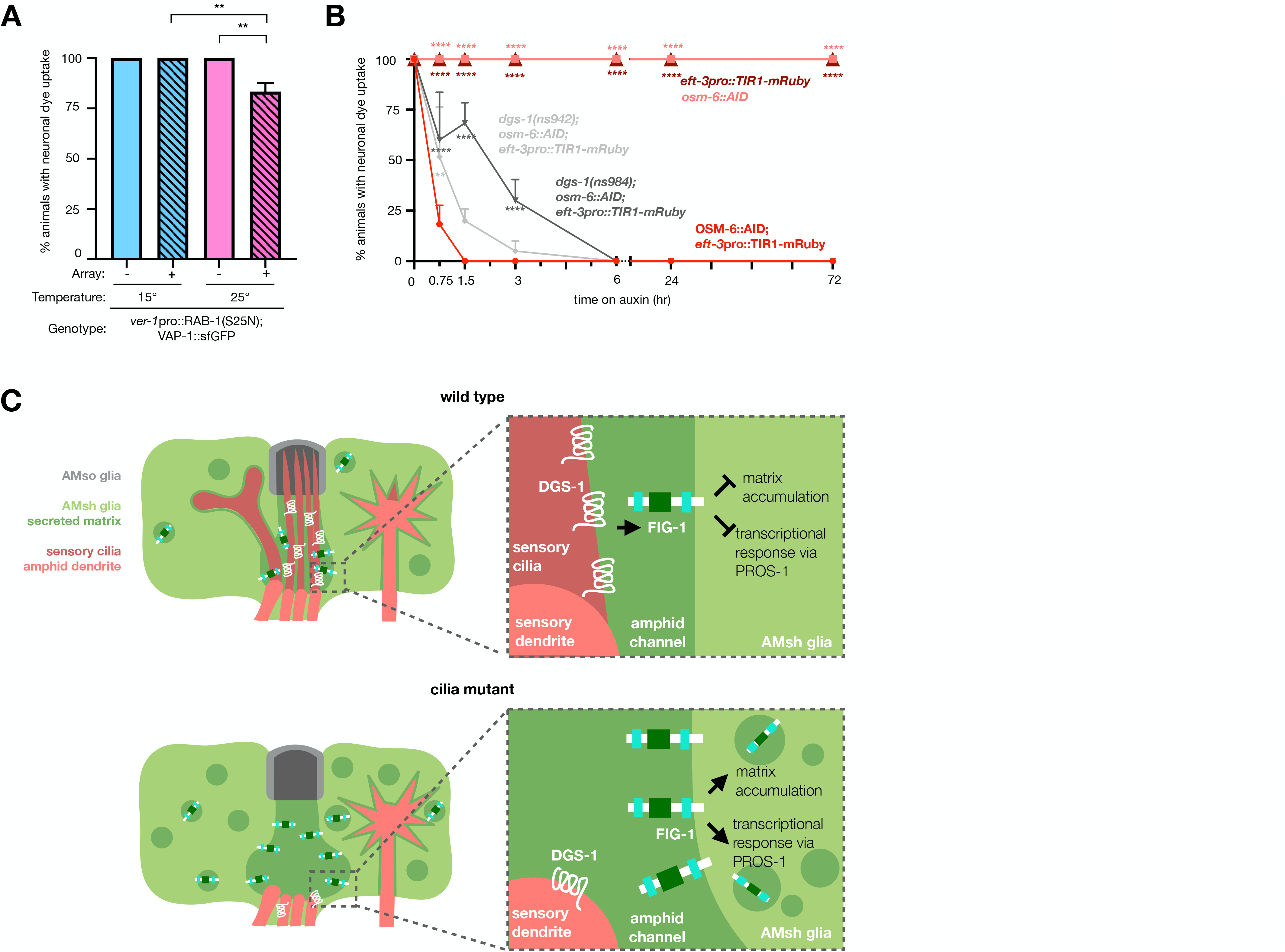
The AMsh glial response is protective to ensheathed dendrites. (**A**) Amphid neuron dye filling in *ver-1*pro::RAB-1(S25N); VAP-1::sfGFP transgenic (array +) and non-transgenic siblings (array −) raised at indicated temperature beginning at the L1 larval stage. Statistical significance calculated using Tukey’s multiple comparison test following 1-way ANOVA. **, P≤0.01. (**B**) Amphid neuron dye filling in animals with indicated genotypes on auxin for indicated times. *n*=3 trials/ condition, 20 animals scored/trial. Data are represented as mean+SEM. Statistical significance calculated using Tukey’s multiple comparison test following 2way ANOVA, shown in relationship to OSM-6::AID; *eft-3*pro::TIR1-mRuby at the same timepoint. (**C**) Model for AMsh glial detection of and response to loss of ensheathed dendrite cilia. Data are represented as mean+SEM.

To test this idea, we used our inducible cilia-disruption system to disrupt cilia in *dgs-1* mutants in which glial secretion is upregulated but dye filling is unaffected. Strikingly, we observed that loss of dye uptake following cilia disruption is significantly delayed in *dgs-1* mutants (**Figure 7B**). While >75% of wild-type animals are dye-filling defective after 45 minutes on auxin, *dgs-1(ns942)* and *dgs-1(ns984)* mutants require at least 1.5 - 3 hours to manifest a similar defect. Importantly, we observe faster dye filling loss in *dgs-1(ns942)* mutants expressing *dgs-1* cDNA under its endogenous promoter (*dgs-1pro*::*dgs-1* cDNA) than in non-transgenic siblings (**Figure S7**).

Our findings, therefore, suggest that AMsh glia secretion is required for normal cilia function, and that the glial response to cilia disruption is protective.

## DISCUSSION

We show here that *C. elegans* AMsh glia detect and mount a protective response to defects in dendritic substructure. While glial responses to axonal structure have been studied extensively, less is known about if or how glia sense the structure of dendrites. Our work now reveals that glia can monitor dendrite structure and respond to changes in dendrite substructure. Our studies uncover key molecular players mediating this glial response. Our findings support a model (**Figure 7C**) in which DGS-1, localizing to cilia of a subset of amphid channel neurons, signals, directly or indirectly, the presence of intact dendrite cilia to the AMsh glia through FIG-1. FIG-1 inhibits AMsh glia matrix accumulation, ensuring optimal levels and composition of AMsh glia-secreted extracellular matrix around cilia. When cilia are damaged, overall DGS-1 levels decrease, and the protein is no longer properly distributed because of cilia structural defects. This attenuates FIG-1 inhibitory activity, resulting in increased secretion from AMsh glia and changes in transcription, including changes in the panel of membrane and secreted proteins released by this cell. The effect of this massive glial response is to protect cilia from damage.

### DGS-1 signals AMsh glia that neuronal cilia are intact

Our findings suggest that DGS-1, a 7-transmembrane domain protein, signals the presence of intact dendrite cilia to surrounding AMsh glia. Many 7-transmembrane-domain proteins function as G protein-coupled receptors (GPCRs), which receive extracellular signals and initiate G protein-dependent signaling cascades. One possibility consistent with our data and with known functions of GPCRs is that DGS-1, constitutively activated by a component of the extracellular matrix mediates a signaling pathway within neurons that releases a neuronal ligand that engages a glial receptor, perhaps FIG-1. Loss of DGS-1 would then block ligand release, triggering the glial response. However, DGS-1 does not have sequence homology to characterized *C. elegans* GPCR families. DGS-1 also does not possess the conserved E/DRY or NPxxY G-protein-coupling motifs found in class A GPCRs, the largest family of GPCRs.^37^ Thus, other models may be useful to consider. If DGS-1 does not function as a GPCR, how might it promote the AMsh glia response? One possibility is that like the multipass-transmembrane proteins MIG-14/Wntless, which promotes Wnt ligand secretion,^38–40^ and CHE-14/ Dispatched, which promote Hedgehog ligand secretion,^41^ DGS-1 may directly promote secretion of a neuronal ligand. Alternatively, DGS-1 may itself be the neuronal signal, and may bind a receptor or other component of the extracellular matrix, perhaps FIG-1, to elicit the AMsh glial responses.

### FIG-1 regulates the AMsh glia cilia damage response

*fig-1b* encodes a protein with a large extracellular domain that regulates AMsh glia secretion. Our structure-function studies of FIG-1b suggest that 2 distinct subdomains within the extracellular region of the protein are important for the glial response to cilia disruption. Interestingly, removal of the short cytoplasmic segment of FIG-1 also increases AMsh glia secretion. This observation suggests that FIG-1 might have intracellular signaling activity, as expected for a signal receptor.

Our studies also reveal that FIG-1 has two separable activities: one involved in promoting associated neuron dye uptake, and another in regulating the AMsh glial response to cilia disruption. Several observations are consistent with the possibility that FIG-1 protein is cleaved, such that the intact transmembrane protein mediates the AMsh glial response, and a cleaved extracellular fragment controls dye uptake. First, both the N-terminal intracellular domain and a proximal extracellular unstructured domain are required for the AMsh glia response but not for dye uptake (**Figures 5A** and **5B**). Second, previous studies from our lab showed that the *fig-1* mutant dye-filling defect can be rescued by expression of the wild-type protein from either neurons or glia.^25^ However, we show here that the AMsh glial response defects in *fig-1* mutants can only be rescued by expression in AMsh glia (**Figure 5F**). Furthermore, we show that a FIG-1::sfGFP fusion protein is detected in two puncta outside of the AMsh glia, localizing to the ends of the two amphid channel cilia bundles, in addition to localizing around cilia (**Figures 5G and 5H**). Localization to these puncta is not consistent with FIG-1 being membrane-bound at this location. Indeed, among AMsh glia-enriched proteins there are at least 3 predicted secreted proteases which could potentially mediate FIG-1 cleavage.^19^ We propose that FIG-1 at the tips of the amphid cilia, which may represent a cleaved version of the extracellular domain, functions in neuronal dye uptake, while intact transmembrane FIG-1 at the base of dendrite cilia, functions in signaling the presence of intact dendrite cilia to AMsh glia. This model is consistent with the observation that *che-2* mutants lose FIG-1::sfGFP localization to cilia tips and display dye filling defects, while *dgs-1* mutants show neither defect.

### Extending the paradigm

Our findings suggest that in the *C. elegans* amphid sensory organ, neurons and the glia that ensheath them engage in a homeostatic interaction tasked with cilia maintenance. Ciliated sensory neuronal endings are found throughout the animal kingdom, and are prevalent not only in nematodes but in insects and vertebrates as well. The overall architecture of these structures is highly conserved. *Drosophila* olfactory sensilla, for example, are composed of sensory neurons whose dendrite cilia are bathed in a specialized matrix directly exposed to the environment, the sensillum lymph, which is secreted by associated glia-like support cells.^42,43^ Similarly, mammalian olfactory sensory neurons are surrounded by glia-like sustentacular cells and their dendrite cilia are bathed in an extracellular matrix secreted largely by nearby Bowman’s gland cells.^44^ Sensory support cells have also received recent attention following the COVID-19 pandemic,^45^ as studies revealed that the genes promoting SARS-CoV2 entry into cells are expressed by sustentacular cells, which have subsequently been implicated as central players in anosmia, or loss of sense of smell, following infection.^46,47^ The proteins that control cilia assembly, including the IFT complex proteins, are highly conserved among these animals,^48^ as is the transcription factor PROS-1/Prospero/Prox1.^19,27–29^ These homologies support the possibility that the cilia-glia interactions we describe here are maintained in additional sensory structures as well.

Ciliated dendrite endings of sensory neurons also resemble dendritic substructures of postsynaptic neurons. Like dendritic spines, cilia are isolated signaling compartments that localize receptors for outside signals. Furthermore, synaptic signals and receptors have molecular similarities to cues received by and receptors localized to cilia.^49^ Thus, it is possible that postsynaptic receptive structures such as dendritic spines communicate with their ensheathing astrocytes to signal their presence. At both sites, glial proteins with thrombospondin domains play important roles: FIG-1 in *C. elegans*, which contains two TSP1 domains controls and responds to cilia, and thrombospondin proteins at mammalian synapses, which regulate synapse formation.^34,50^ It is also of note that a subfamily of mammalian adhesion GPCRs, the brain-specific angiogenesis inhibitors (BAIs), resemble a fusion of FIG-1 and DGS-1 in domain architecture and topology and are subsequently cleaved into distinct peptides. BAIs are expressed in mammalian neurons and regulate synaptogenesis and dendritic spine formation.^51^ Might these protein represent a fusion between DGS-1 and FIG-1 functions? While our attempts to rescue *C. elegans dgs-1*; *fig-1* double mutants with a FIG-1/ DGS-1 fusion have not been successful to date, the possibility warrants continued pursuit.

## ACKNOWLEDGEMENTS

We thank Piali Sengupta and Shohei Mitani for strains; Aakanksha Singhvi and Piali Sengupta for plasmids; members of the Shaham lab for useful discussions and reagents; and the Rockefeller University Flow Cytometry and Genomics Resource Centers for technical support. Some strains were provide by the CGC, which is funded by the NIH office of Research Infrastructure Programs (P40 OD010440). K.C.V. was a Kavli Neural Systems Institute postdoctoral fellow. Research reported in this publication was supported by the National Institute of Neurological Disorders and Stroke of the National Institutes of Health under Award Number F32NS105322 to K.C.V. and R35NS105094 to S.S. The content is solely the responsibility of the authors and does not necessarily represent the official views of the National Institutes of Health.

## AUTHOR CONTRIBUTIONS

Conceptualization, K.C.V and S.S.; Methodology, K.C.V., B.M.H., L.L., A.F, and S.S.; Software, Y.Liang, Formal Analysis, K.C.V. Investigation, K.C.V, B.M.H., L.L., A,F., and Y. Lu; Writing-Original Draft, K.C.V. Writing-Review & Editing, K.C.V., B.M.H, L.L, A.F, and S.S.; Funding Acquisition, K.C.V. and S.S.

## MATERIALS AND METHODS

### Caenorhabditis elegans strains and handling

Experiments were performed in synchronized populations of one day old hermaphrodites, unless otherwise noted. Animals were grown on nematode growth media plates seeded with *E. coli* strain OP50 as a food source, with the exception of animals used for RNA-seq experiments which were grown in liquid culture with *E. coli* strain HB101 as a food source. Strains were maintained using standard methods.^60^ *C. elegans* Bristol strain N2 (RRID:WB-STRAIN:WBStrain00000001) was used as wild type.

### CRISPR/Cas9 genome editing

*vap-1(ns831[vap-1*::sfGFP]) was generated using plasmid-based methods.^61^ An sgRNA guide sequence targeting the 3’ end of *vap-1* (5’-gttgcatagaaaatttacta-3’) was cloned into pDD162, creating plasmid pKV2. A homologous recombination template plasmid, pKV3, was cloned, containing the following elements: (1) 1.5kb *vap-1* 5’ homology arm of *vap-1* coding sequence, (2) an in frame flexible linker coding for amino acid sequence: PDPRDWPKDRK, (3) an artificial intron containing *unc-119*(+) rescuing sequence in the antisense direction flanked by LoxP sites, (3) additional in frame flexible linker coding for amino acid sequence: EDPWRVP, (4) in frame sfGFP coding sequence, (5) 1.5kb *vap-1* 3’ homology arm of intergenic sequence, and (6) plasmid backbone from pPD95.75. Note that a silent mutation was introduced in the final codon of the sfGFP sequence to disrupt a potential PAM site immediately following the sgRNA guide target site. pKV2 and pKV3 were injected along with red fluorescent co-injection markers into strain DP38. Progeny were screened for rescue of uncoordinated phenotype without expression of red co-injection markers and genotyped for insertion of desired sequence into the *vap-1* locus. Once insertion of the desired sequence was verified, animals were injected with pPD104 to remove *unc-119*(+) rescuing sequence and backcrossed with N2 to remove the *unc-119(ed3)*III mutation, creating strain OS11927.

*dgs-1(ns984)* was generated by imprecise insertion via injection of Cas9, tracrRNA, and crRNA from IDT.^62^

The following alleles were generated by SUNY biotech:

- *osm-6(syb2906[osm-6::gfp-aid])*V - C-terminal knock-in of coding sequences of: (1) linker sequence: GASGASGAS, (2) GFP, and (3) AID
- *osm-6(syb2906 syb4401[osm-6::gfp-aid])*V - precise deletion of GFP sequence from *osm-6(syb2906)*
- *dgs-1(ns942 syb3113)*IV - precise nucleotide replacement, reverting *dgs-1(ns942[G58>E])* allele to wild type, E58>G
- *fig-1(syb6983[fig-1Δcoding sequence])V -* precise deletion of *fig-1* coding sequence
- *fig-1(syb5898[fig-1ΔSS/TMD])V -* precise deletion of *fig-1b* transmembrane domain (FIG-1B amino acids 23-51)
- *fig-1(syb7619[fig-1ΔC-terminus])V -* precise deletion of *fig-1* C-terminus (FIG-1B amino acids 2,874-2,898)
- *fig-1(syb7600[fig-1ΔN-terminus])V -* precise deletion of *fig-1b* N-terminus (FIG-1B amino acids 2-22)
- *fig-1(syb7606[fig-1Δunstructured])V -* precise deletion of *fig-1b* unstructured extracellular region (FIG-1B amino acids 115-754)
- *fig-1(syb6326[fig-1Δtsp C])V -* precise deletion of *fig-1* C-terminal thrombospondin type 1 repeat (FIG-1B amino acids 2,823-2,873)
- *fig-1(syb6326 syb6968[fig-1Δtsp N + tsp C])V -* precise deletion of *fig-1* N-terminal and C-terminal thrombospondin type 1 repeats (FIG-1B amino acids 62-114 & 2,823-2,873)
- *fig-1(syb7051[fig-1Δisoform a])V -* precise deletion of *fig-1b* intron 19 containing the first exon of *fig-1a*
- *fig-1(syb5954[fig-1Δtsp N])V -* precise deletion of *fig-1b* N-terminal thrombospondin type 1 repeat (FIG-1B amino acids 62-114)
- *fig-1(syb6039[fig-1ΔC6])V -* precise deletion of *fig-1b* region containing 17 C6 repeats (FIG-1B amino acids 755-2,788)
- *fig-1(syb7231[fig-1::sfGFP C-terminal])V -* C-terminal knock-in of coding sequences for: (1) flexible linker with amino acid sequence: PDPRDWPKDPK (2) sfGFP
- *fig-1(syb7028[fig-1::sfGFP internal)V* knock-in of coding sequences for the following after FIG-1B amino acid 754: (1) flexible linker with amino acid sequence: PDPRDWPKDPK, (2) sfGFP, (3) flexible linker with amino acid sequence: GGSGGGSGGGSG

### Generation of recombinant DNA via PCR fusion

Several fluorescent reporters and rescuing DNA sequences were generated using a PCR fusion approach.^63^ First, two fragments were amplified via PCR: (1) promoter sequences containing 24-29bp of homology region to the 5’ sequence of fragment 2, added via the reverse PCR primer, and (2) coding sequence of fluorescent report or rescuing cDNA and UTRs. The two fragments were then fused via PCR and amplified using internal nested primers. *F16F9.3*pro::myr-mKate2 was created by fusing: (1) 2,056bp *F16F9.3* promoter region and (2) coding sequence for myr-mKate2 followed by the *unc-54* 3’UTR. *dgs-1pro*::GFP was created fusing: (1) an intergenic region upstream of *dgs-1* and (2) GFP followed by the *unc-54* 3’UTR. *dgs-1*pro::*dgs-1* cDNA was created by fusing: (1) an intergenic region upstream of *dgs-1* and (2) *dgs-1* cDNA with endogenous 5’ and 3’ UTRs. *dyf-11*pro::*dgs-1* cDNA was created by fusing: (1) a 1,864bp *dyf-11*pro and (2) *dgs-1* cDNA with endogenous 5’ and 3’ UTRs.

### Plasmid construction

Plasmids were constructed using Gibson cloning.

Cell-specific TIR1-mRuby plasmids were created by subcloning TIR1-mRuby followed by the *unc-54* 3’UTR amplified from pLZ31^30^ (fragment) into plasmids containing the following cell-specific promoters (vectors): (1) a 1,864bp *dyf-11*pro (pKV13), (2) a 2,056bp *F16F9.3*pro (pKV14), (3) a 3.2kb *srh-142*pro (pKV16) (4) a 1.0kb *srg-47*pro (pKV17), (4) the *dgs-1*pro (pKV15).

DGS-1::mKate2 (pKV12) was created by Gibson assembly of 3 fragments: (1) a vector containing the *dgs-1* promoter, endogenous 5’ UTR, and *dgs-1* cDNA to the center of the 3^rd^ intracellular domain, (2) mKate2 flanked by flexible linkers (PDPRDWPKDPK/GGSGGGSGGGSG), and (3) the remaining *dgs-1* cDNA followed by the endogenous 3’UTR.

*fig-1*pro::*fig-1*ΔC6 cDNA (pKV34) was created by inserting *fig-1*b cDNA in two fragments (1) amino acids 1-754 and (2) amino acids 2,789-stop into a vector containing a 1,872bp *fig-1* promoter. *F16F9.3*pro:: *fig-1*ΔC6 cDNA (pKV35) and *dyf-11*pro:: *fig-1*ΔC6 cDNA (pKV36) were generated by replacing the *fig-1*pro in pKV34 with: (1) a 2,056bp *F16F9.3* or (2) 1,864bp *dyf-11* promoter, respectively.

### Germline transformation and integration

Plasmid mixes containing the plasmid(s) of interest, co-injection markers, and pBluescript were injected into the gonads of young adult hermaphrodites at a total of 100 ng/ul. Injected animals were singled onto NGM plates and allowed to grow for one generation. Transformed animals were screened for the expression of fluorescent co-injection markers, singled, and screened for stable inheritance of the extrachromosomal array. Only distinct F1s or lines from different P0 injected hermaphrodites were considered independent.

Integrated transgenic constructs (*nsIs971* and *nsIs972*) were generated by exposure to 33.4 μg/mL trioxsalen and UV irradiation using a Stratagene Stratalinker UV 2400 Crosslinker (360 μJ/cm2 x100).^64^

### Microscopy

Animals were anesthetized using 50mM NaN_3_ and mounted on 2% agarose pads on glass slides. Widefield z-stacks (0.3-1μm thick) were taken using a Zeiss compound microscope (Axio Imager M2) using a 63X objective controlled by MicroManager software.^65^ Confocal z-stacks (0.3-1μm thick) were taken using a Zeiss LSM900 using a 63X objective controlled by ZenBlue software. ImageJ software was used to produce maximum projections of z-stack images.

### VAP-1::sfGFP accumulation scoring

Animals were anesthetized on ice for ~10-60 minutes prior to observation with a fluorescence dissecting microscope equipped with a 2X objective (Leica). Animals with obvious GFP accumulation at the nose tip were scored as strong, animals in which it was unclear if GFP accumulation was in excess of wild type were scored as weak, and animals with no visible GFP fluorescence were scored as none.

### Cell isolation and FACS analysis

~2 million synchronized larvae expressing *F16F9.3*pro::dsRed in wild type or *che-2(e1033)* background (OS4079 and OS11549) were grown from L1 arrest in S-basal liquid culture containing *E. coli* HB101 at 20°C, shaking. After 36-42 hours, late-stage larvae (L3 and L4) were pelleted by centrifugation (2 min, 1,300 rpm) and subsequently washed ten times with M9 to remove excess bacteria. Each wash consisted of a brief (10 second, 1300 rpm) centrifugation, such that most animals were pelleted, but bacteria remained in suspension. Animals were then dissociated using SDS-DTT (0.25% SDS; 200 mM DTT; 20 mM HEPES, pH 8.0; 3% sucrose) and Pronase E (15 mg/ml), as previously described.^66^ SDS-DTT was added at a 2:1 ratio the volume of packed animal pellet, followed by 4 min incubation on ice. After washing, 4:1 ratio of Pronase E was added to the packed animals’ pellet and animals were incubated rotating at 20°C for 5 min, followed by 12 min of gentle homogenization (2mL dounce homogenizer, pestle clearance 0.0005-0.0025 inches). After washes with ice-cold egg buffer (1.18 M NaCl; 480 mM KCl; 20 mM CaCl_2_; 20 mM MgCl_2_; 250 mM HEPES, pH 7.3) to remove Pronase E, cells were filtered through a 5μM filter to remove undigested animal fragments, and immediately sorted by FACS.

AMsh glia cell sorting was performed using a BD FACS Aria sorter (Rockefeller University Flow Cytometry Resource Center), with egg buffer as the sheath buffer to preserve cell viability. Dead cell exclusion was carried out using DAPI, while DRAQ5 was used to distinguish nucleated cells from non-nucleated cell fragments. Gates for size and granularity were adjusted to exclude cell aggregates and debris. Gates for fluorescence were established using wild type non-fluorescent animals. 168,356-240,125 dsRed-positive events were sorted per replicate, which represented 0.1−0.4% of total events (after scatter exclusion), which is roughly the expected labeled-cell frequency in the animal (~0.21%). dsRed-negative events from the same gates of size and granularity, representing all other cell types, were also sorted for comparison. Cells were sorted directly into TRIzol LS at a ratio 3:1 (TRIzol to cell volume).

### RNA isolation and sequencing

RNA was extracted from Trizol LS-treated cells by phase separation, following the manufacturers guidelines. RNA was purified using PicoPure RNA isolation kit. 0.4-5ng purified total RNA was obtained per sample. All subsequent steps were performed by the Rockefeller University Genomics Resource Center. RNA quality was verified using an Agilent Bioanalyzer to ensure sample degradation had not occurred. mRNA amplification and cDNA preparation were performed using the SMARTer mRNA amplification kit. Labelled samples were sequenced using an Illumina HiSeq 2000 sequencer and standard Illumina sequencing primers.

### RNA-seq quality assessment

Fastq files were generated with CASAVA v1.8.2, and examined using FASTQC for sequence quality. Reads were aligned to *C. elegans* WS262 genome release (https://downloads.wormbase.org/releases/WS262/species/c_elegans/PRJNA13758/) using the STAR v2.3 aligner with parameters (--out-FilterMultimapNmax 10 --outFilterMultimapScoreRange 1). The alignment results were evaluated using RNA-SeQC v1.17 to make sure all samples had a consistent alignment rate and no obvious 5′ or 3′ bias. Aligned reads were summarized through featureCounts with gene models from Ensemble (Caenorhabditis_elegans. WBcel235.77.gtf) at gene level unstranded. Specifically, the uniquely mapped reads (NH “tag” in bam file) that overlapped with an exon (feature) by at least 1 bp on either strand were counted and then the counts of all exons annotated to an Ensemble gene (meta features) were summed into a single number. rRNA genes, mitochondrial genes and genes with length <40 bp were excluded from downstream analysis.

### RNA-seq differential gene expression analysis

Experiments were performed with 3 independent replicates. DESeq2 was applied to normalize count matrix and to perform differential gene-expression analysis, comparing: (1) RNA counts derived from the AMsh glia cells (dsRed positive) to RNA counts that were derived from all other *C. elegans* cells (dsRed negative) from both wild type and cilia mutant *[che-2(e1033)]*, and (2) AMsh glia from wild type to AMsh glia from cilia mutant, using negative binomial distribution.^67^ To identify transcripts up- or down-regulated in cilia mutant compared to wild type AMsh glia, we used a fold-change cut offs of >2 and <0.5, respectively, and an adjusted p-value threshold of <0.1.

### Inducible cilia disruption

Exposure of *C. elegans* to the synthetic auxin analog K-NAA (1-napthaleneacetic Acid Potassium Salt), results in ubiquitination and subsequent proteasomal degradation of auxin inducible degron (AID)-tagged proteins in the presence of transgenically provided TIR1, the substrate recognition component of the E3 ubiquitin ligase complex.^30,68^ Strains carrying *osm-6::gfp::aid [osm-6(syb2906)]* or *osm-6::aid (osm-6(syb2906 syb4401)* were crossed with a strain ubiquitously expressing TIR1 tagged with mRuby (*ieSi57 [eft-3pro::TIR1-mRuby*]) or injected with constructs expressing TIR1-mRuby under cell specific promoters. K-NAA was dissolved in sterile water to prepare a 200mM stock solution. For 72 hour auxin treatment, OP50-seeded NGM plates were pre-coated with K-NAA to a final concentration of 4mM and synchronized populations of animals were transferred to K-NAA plates. For all other timepoints, 200mM K-NAA was added directly to animals on seeded plates and allowed to dry before the remaining incubation.

### Neuronal Dye Filling

5mg/mL DiI (1,1’-dioctadecyl-3,3,3’3’-tetramethylindocarbocyanine perchlorate) and 2mg/mL DiO (3,3′-Dioctadecyloxacarbocyanine perchlorate) stock solutions were made in N,N-dimethylformamide. Animals were incubated with dye solution, diluted in M9 to a final concentration of 5μg/ml for DiI or 8μg/mL for DiO, for (1) 15-60 minutes for routine dye filling assays or (2) 5 minutes for dye filling time course experiments. Animals were then washed twice in M9, and transferred to seeded plates at least 10 minutes to remove excess dye prior to scoring. Dye filling was scored as complete, partial, or none, as compared to wild type using fluorescence dissecting microscopes. For dye filling time course experiments, animals were anesthetized on ice for ~10-60 minutes prior to scoring and dye filling was scored in the cell bodies of neurons.

### Mutagenesis and mutant identification

OS12214 animals were mutagenized using 75 mM ethylmethanesulfonate for 4h at 20 °C. ~10,000 F2 progeny were screened for VAP-1::sfGFP accumulation and animals with strong accumulation were transferred to individual plates. A secondary screen was performed to exclude isolates with neuronal dye filling defects, indicating defects in dendrite cilia.

### SNP mapping

*ns942* mutants were crossed to CB4856 Hawaiian males. F2 animals with VAP-1::sfGFP accumulation were isolated, and progeny were lysed and genotyped for 3 SNPs between N2 and Hawaiian backgrounds on each chromosome. Additional SNPs were used to map *ns942* to a ~4.5 map unit segment of chromosome IV.

### Whole genome sequencing

*ns942* and parental OS12214 animals were grown on *E. coli* strain OP50 until bacteria were depleted, harvested in M9, and resuspended in 0.5mL TEN buffer (20 mm Tris pH 7.5, 0.5 M EDTA, 100 mM NaCl), pH 7.5. SDS (0.5%), proteinase K (0.1 mg/mL), and β-mercaptoethanol (0.2%) were added. The lysis reaction was incubated overnight in a shaking thermocycler at 56°C. Phenol/chloroform was added and phase-separated by spinning in a phase-lock tube. The aqueous phase was transferred to 200 proof EtOH. The resulting DNA clot was washed in 70% EtOH, dried, and resuspended in TEN. After rehydration, 0.3 μL of 100 mg/mL RNAse A was added and incubated at 37°C for 2 h. Phenol/chloroform extraction was performed again and DNA was rehydrated with EB buffer. The sample was then examined using a nanodrop spectrophotometer and run on a 1% agarose gel to confirm that there was no RNA contamination or DNA degradation, respectively. NextSeq Mid Output 2 x 150 sequencing was performed using Standard Illumina Sequencing primers for gDNA-seq application.

### Electron Microscopy

Synchronized young adult animals were fixed, stained, embedded in resin, and sectioned using standard methods.^69^ For OSM-6::AID; *eft-3*pro::TIR1-mRuby; *vap-1(ns831)* animals treated with auxin for 3 hours, after 1.5 hours of auxin treatment, an abbreviated dye filling assay was performed in the presence of 4mM K-NAA, and dye filling defective animals were selected for imaging after 3 hours auxin treatment total. Serial images were acquired by using a Titan Themis 200 kV transmission electron microscope with Cs Image Corrector. Image processing and analysis were performed using ImageJ and IMOD software.

### AMsh-specific secretion block experiments

*ver-1*pro::RAB-1(S25N); VAP-1::sfGFP (OS12934) bleach synchronized L1 larvae were grown to young adulthood at either 15C (4 days) or 25C (2 days), when DiO dye filling assay was performed.

## QUANTIFICATION AND STATISTICAL ANALYSIS

All fluorescence microscopy quantifications were performed in ImageJ.^59^ Control and experimental animals imaged during the same imaging session with all acquisition parameters maintained constant between the conditions. For quantification of amphid fluorescence, a trapezoidal region of interest (ROI) was drawn in the DIC channel, with the nose tip as the anterior boundary and the buccal cavity as the posterior boundary. For quantification of cell body fluorescence, a rectangular ROI wad drawn around the diameter of the animal where cell body fluorescence was observed. For quantification of puncta/ animal, puncta were followed and counted through z-stacks through entire worm, comparing to relevant ROI for size comparison as noted. Mean, background subtracted fluorescence is reported. If normalized, the mean of the wild type condition was used for each experimental imaging session was used for normalization.

Prism software was used for statistical analysis, with statistical tests noted in figure legends. For gene list overlap statistics, normal approximation to the hypergeometric probability was calculated (http://nemates.org/MA/progs/overlap_stats.html; Jim Lund, personal communication).

---

**Figure S1.**
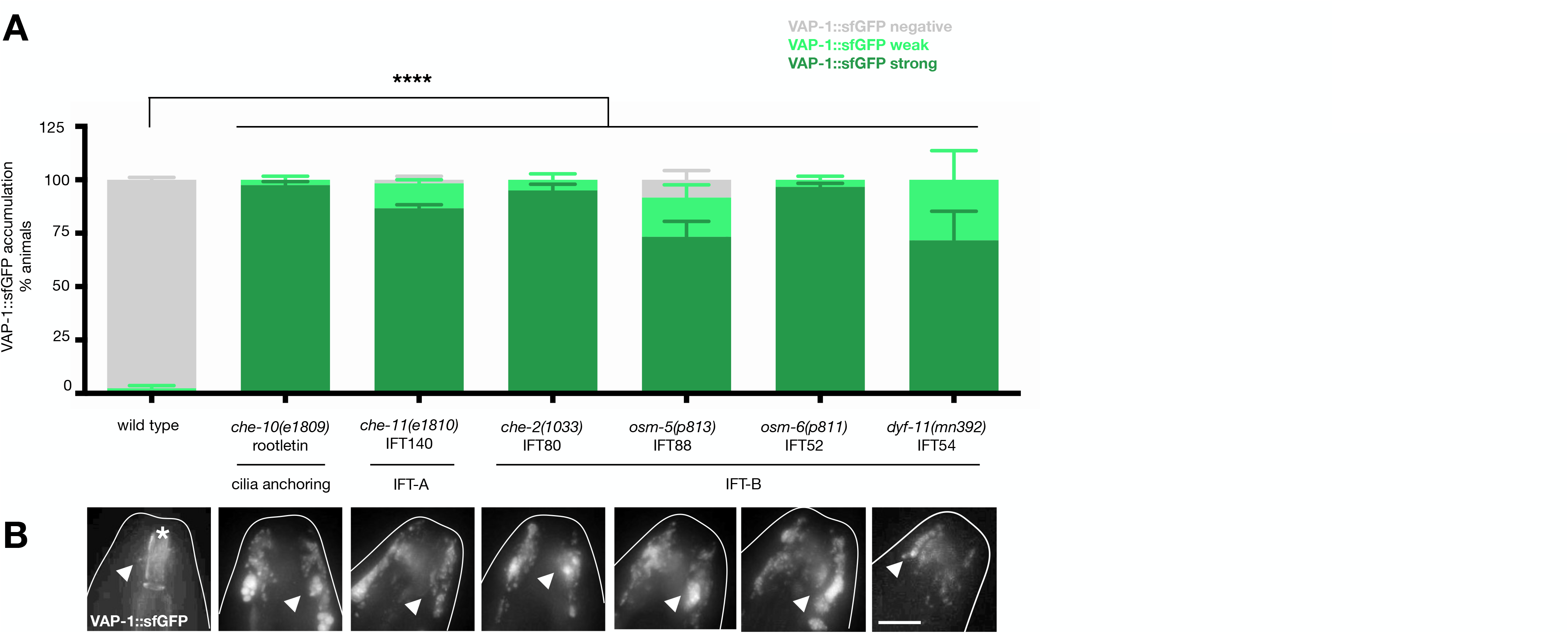
AMsh glia ensheathing dendrites with diverse mutations in sensory cilia genes respond by accumulating secreted matrix. VAP-1::sfGFP accumulation scoring (**A**) and representative brightness and contrast matched images of VAP-1::sfGFP (**B**) of wild type and indicated cilia mutant animals. Arrowheads,VAP-1::sfGFP puncta. Asterisk, anterior buccal cavity auto-fluorescence. Data are represented as mean+SEM of n≥3, 20 animals/ trial. Statistical significance calculated using Dunnett’s multiple comparison test following one-way ANOVA. ****, P≤0.0001. Images are maximum intensity projections of widefield z-stacks. Scalebar= 10μm.

**Figure S2.**
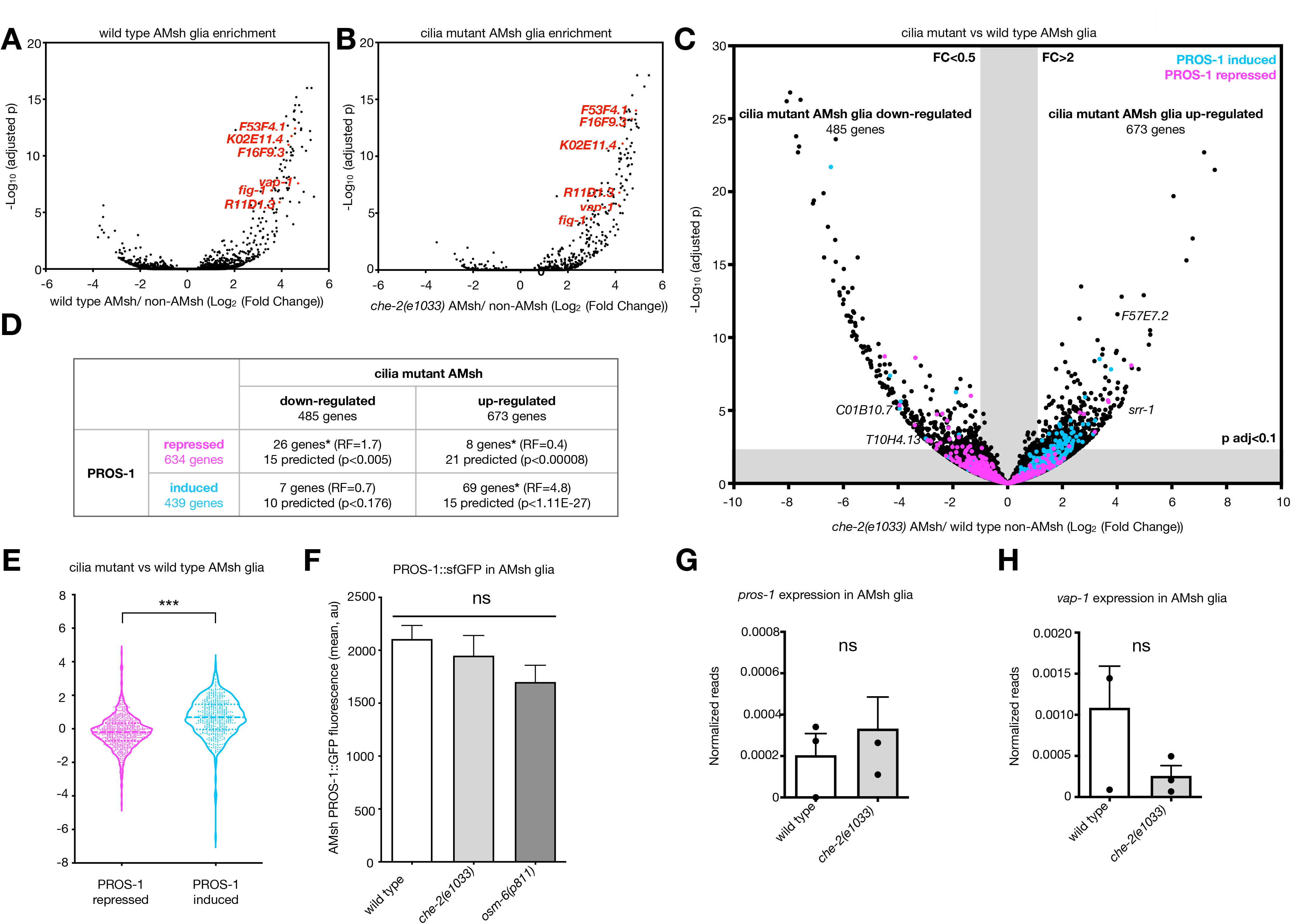
AMsh glia transcriptionally respond to defects in ensheathed dendrite cilia, correlating with increased PROS-1 activity. Volcano plots of AMsh enriched genes (reads in purified dsRed + AMsh glia compared to dsRed-cells) from wild type (**A**) and cilia mutant [*che-2(e1033)*] animals (**B**). Known AMsh glia-enriched genes are highlighted in red. (**C**) Volcano plot of gene expression changes between purified wild type and cilia mutant [*che-2(e1033)*] AMsh glia. PROS-1 induced and repressed genes are labeled in cyan and magenta, respectively.^19^ (**D**) Table comparing genes down- and up-regulated in cilia mutant versus wild type AMsh glia and genes regulated by PROS-1. Total number of genes in each group and % of total protein-coding genes in the *C. elegans* genome is listed for each group. Normal approximation to the hypergeometric probability was calculated. *, P≤0.005. (**E**) Violin plot of PROS-1 induced and repressed genes plotted by enrichment in cilia mutant versus wild type AMsh glia. Statistical significance calculated using unpaired t test. ***, P≤0.001. (**F**) Quantification of PROS-1::GFP fluorescence in wild type and two cilia mutants: *che-2(e1033)* and *osm-6(p811)*. (**G**) *pros-1* and (**H**) *vap-1* expression (normalized RNA-seq reads) in AMsh glia FACS-purified from wild type and cilia mutant [*che-2(e1033)*] animals. Data are represented as mean+SEM. Statistical significance calculated by Dunnett’s multiple comparison test following ordinary one-way ANOVA.

**Figure S3.**
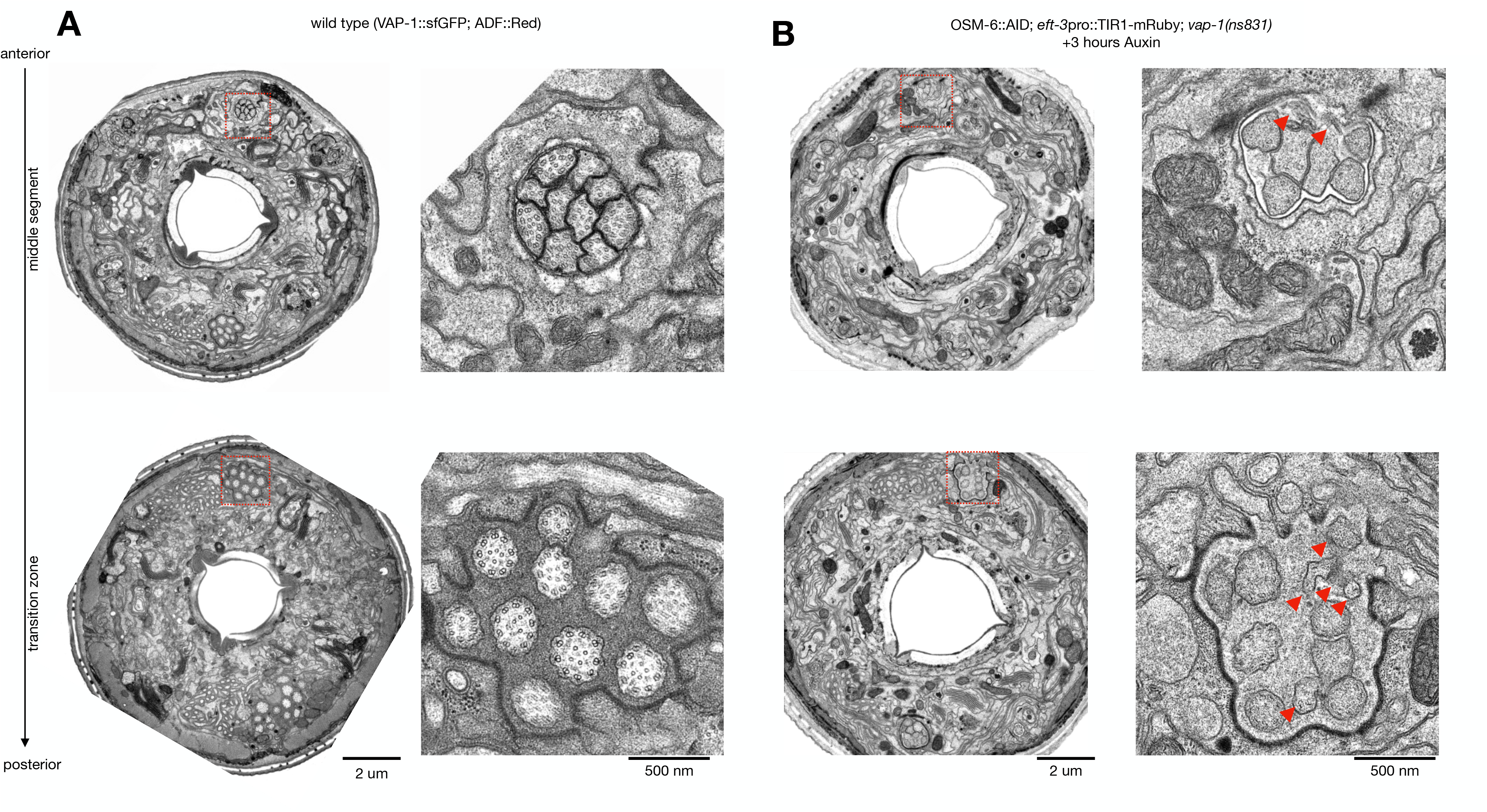
Inducible OSM-6 degradation causes acute cilia disruption. TEM cross sections of through (**A**) an untreated wild-type animal (VAP-1::sfGFP; (*srh-142pro::dsRed*)) and, (**B**) a dye filling defective OSM-6::AID; *eft-3*pro::TIR1-mRuby; *vap-1(ns831)* animal following 3 hours of auxin treatment. Whole animal cross sections are shown on the left with enlarged insets of channel cilia at higher resolution shown on the right. Both the middle segment (anterior) and transition zone (posterior) regions of the cilia are shown for each animal. Arrows, abnormal cilia morphology.

**Figure S4.**
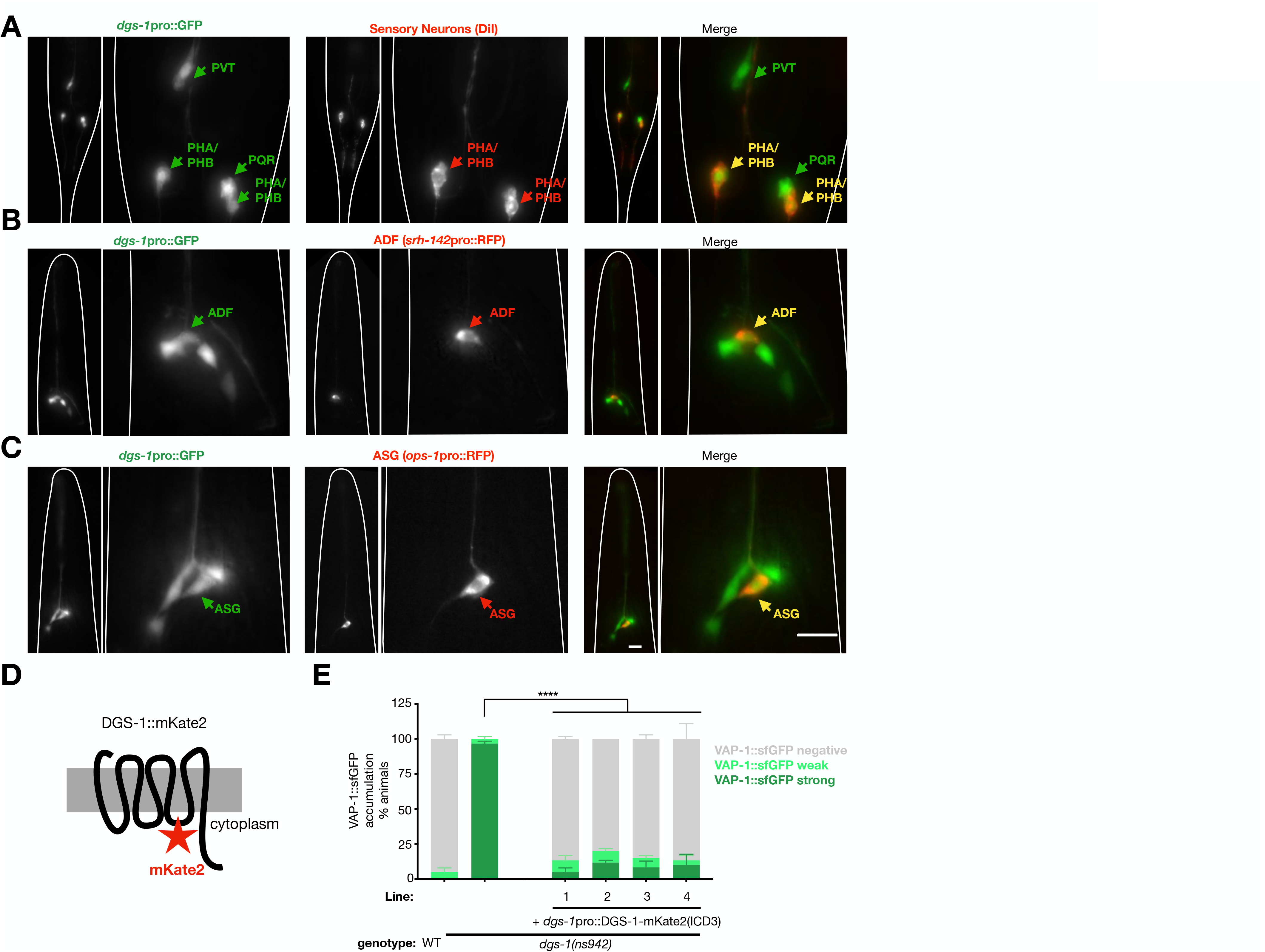
*dgs-1* is expressed in phasmid, ADF, and ASG sensory neurons and is functional when tagged in the 3^rd^ intracellular domain. Representative images of animals expressing *dgs-1*pro::GFP co-labeled (**A**) with DiI (subset of sensory neurons), (**B**) an ADF neuron reporter (*srh-142*pro::RFP), or (**C**) an ASG neuron reporter (*ops-1*pro::RFP). Scale bars=10μm. (**D**) DGS-1::mKate2 predicted topology. (**E**) VAP-1::sfGFP accumulation in wild type, *dgs-1(ns942)*, and *dgs-1(ns942)* expressing DGS-1 fused to mKate2 in the 3^rd^ intracellular domain. Data represented as mean+SEM. Statistical significance calculated using % animals with strong VAP-1::sfGFP accumulation using Dunnett’s multiple comparison test following ordinary one-way ANOVA. ****, P≤0.0001.

**Figure S5.**
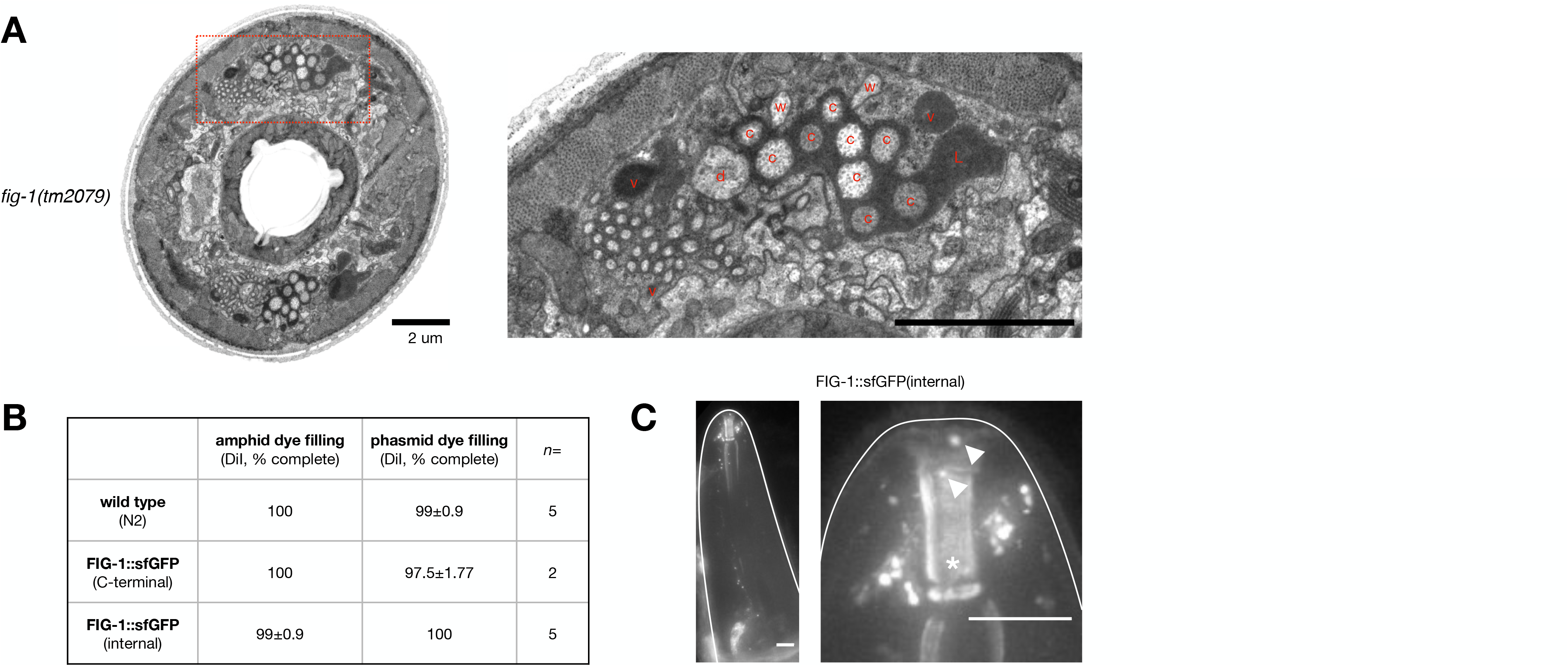
*fig-1(tm2079)* mutants accumulate AMsh secreted matrix, FIG-1::sfGFP fusions do not cause dye filling defects, and FIG-1::sfGFP(internal) also accumulates in anterior puncta. (**A**) Electron micrograph cross sections through *fig-1(tm2079)* mutant animals, with enlarged inset of the right amphid shown in red dotted lines. L, amphid channel lobes. v, matrix filled AMsh vesicle. c, channel cilia. d, dendrite. w, wing cilia. Scale bars=2μm. Samples originally prepared for previous publication.^25^ (**B**) Dye filling data of FIG-1::sfGFP strains. *n* indicated in table, 20 animals/trial. Data represented by mean+SEM. (**C**) Widefield z-stack of FIG-1::sfGFP(internal) animal. Arrowheads, anterior FIG-1::sfGFP puncta, Asterisk, posterior buccal cavity auto-fluorescence. Scale bars=10μm.

**Figure S6.**
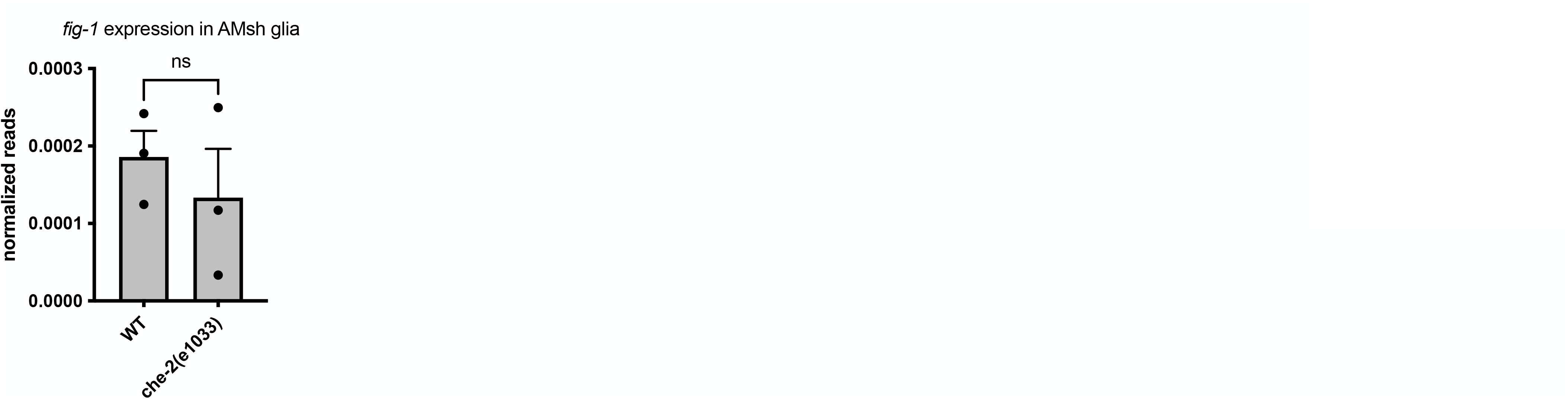
*fig-1* transcription is not changed in cilia mutants. *fig-1* expression (normalized RNA-seq reads) in AMsh glia FACS-purified from wild type and cilia mutant [*che-2(e1033)*] animals. Data represented by mean+ SEM. Statistical significance calculated by unpaired t test. ns, P>0.05.

**Figure S7.**
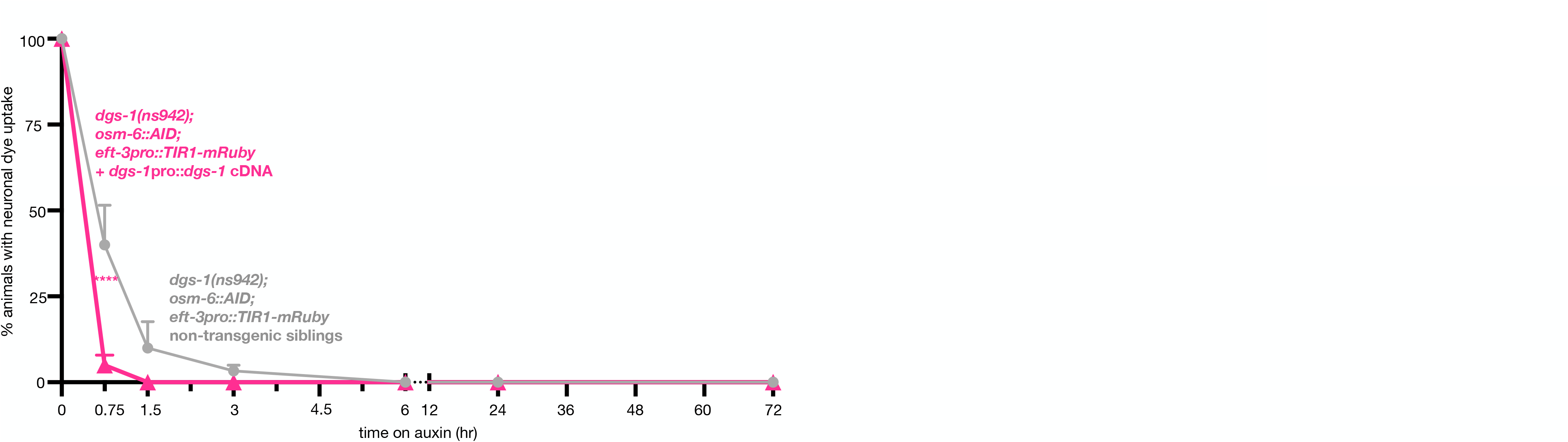
Expression of *dgs-1* cDNA prevents protective effect of the glial response to loss of dendrite cilia in *dgs-1* mutants. Amphid dye filling in the neurons of animals with indicated genotypes on auxin for indicated time. *n*=3 trials/condition, 20 animals/trial. Data are represented as mean+SEM. Statistical significance calculated using Šídák’s multiple comparison test following 2-way ANOVA and shown in relationship to non-transgenic siblings at the same timepoint. ****, P≤0.0001.

